# Prenatal arsenic exposure stymies gut butyrate production and enhances gut permeability in post natal life even in absence of arsenic deftly through miR122-Occludin pathway

**DOI:** 10.1101/2022.06.13.496028

**Authors:** Mainak Chakraborty, Anupam Gautam, Oishika Das, Aaheli Masid, Moumita Bhaumik

**Author notes:** Correspondence: Moumita Bhaumik, ICMR-National Institute of Cholera and Enteric Diseases, P-33 C.I. T Road, Beleghata, Kolkata 700010, West Bengal, India.

## Abstract

This discourse attempts to capture a few important dimensions of gut physiology like microbial homeostasis, short chain fatty acid (SCFA) production, occludin expression and gut permeability in post-natal life of mice those received arsenic only during pre-natal life (pAs-mice). The pAs-mice showed a striking reduction in Firmicutes to Bacteroidetes (F/B) ratio coupled with decrease in tight junction protein, occludin resulting in increase in gut permeability, increased infiltration of inflammatory cells in the colon and decrease in common SCFAs in which butyrate reduction was quite prominent in fecal samples as compared to normal control. The above phenotypes of pAs-mice were mostly reversed by supplementing butyrate with food. The talismanic ability of butyrate in enhancing occludin expression, in particular, was dissected further. As miR122 causes degradation of Occludin mRNA, we transiently overexpressed miR122 by injecting appropriate plasmids and showed reversal of butyrate effects in pAs-mice. Thus, pre-natal arsenic exposure orchestrates variety of effects by decreasing in butyrate in pAs-mice leading to increased permeability due to reduced occludin expression. Our research adds a new dimension to our understanding that pre-natal arsenic exposure imprints in post-natal life while there was no further arsenic exposure.

**Highlights:**

- Prenatal Arsenic exposure decreases prevalence of butyrate producing bacteria and butyrate production in gut.
- Lack of butyrate production in the gut is responsible for increased permeability and decreased occludin expression.
- Oral supplementation with butyrate reverses the prenatal arsenic induced changes in the gut.
- Butyrate increases Occludin gene expression by downregulating miR122 in the gut.

## 1. Introduction

Arsenic is a highly potent environmental toxin affecting more than 200 million people globally where south east Asian countries including India, Bangladesh, Pakistan, Nepal and China are largely affected (Mukherjee et al. 2006). As arsenic is readily absorbed *in utero* the intake of arsenic by children (per unit body mass) is higher than that of adults increasing the risk of infant mortality, health risks and impaired intellectual development with associated impacts later in life (Donohoe et al. 2012), (Cherry et al. 2010), (Myers et al. 2010). Earlier we have shown that prenatal arsenic exposure causes compromised immune response in juveniles which lead to susceptibility to pathogen infection (Chakraborty and Bhaumik 2020). Recent reports showed that environmentally relevant level of arsenic exposure in mice causes gut microbial dysbiosis (Chi et al. 2017). Although the children in arsenic affected areas showed altered gut microbial composition (Dong et al. 2017), oddly enough there are no reports till date on how *in utero* arsenic exposure may change the gut microbiome and its consequences on gut function, if any, in later life.

Maternal–offspring microbiota exchanges play a significant role in the development and maturation of the neonatal microbiome (Dong et al. 2015). Epidemiological studies also indicate antibiotic usage during pregnancy has also been linked to an increased incidence of juvenile obesity (Mor et al. 2015) and asthma (Stensballe et al. 2013). It has also been reported that the children born to mothers suffering from ulcerative colitis whose disease was active during pregnancy have a higher risk of developing childhood illnesses (Hashash and Kane 2015). Recent reports showed that arsenic exposure in early life alters the gut microbiome during the critical window of infant development (Hoen et al. 2018), (Dong et al. 2017). These observations festoon an impending issue which nudge us to investigate impact of prenatal arsenic exposure on the gut physiology in post-natal life. Germane with the idea that mother’s microbiome plays a role in infant’s gut microbial establishment; it is plausible that subtle alterations in microbiota in early life may act as a vulnerability factor, impacting on disturbances in gut physiological functions which may lead to disorders in adult hood.

Short chain fatty acids (SCFA), an important metabolite produced by the gut microbes after fermentation of dietary fibres are significantly decreased in arsenic exposure (Chi et al. 2017). Butyrate, one of the important SCFA controls pleotropic functions in the body including development of the immune system (Yip et al. 2021), (Schulthess et al. 2019). Butyrate can down-regulate inflammation by inhibiting the growth of pathobionts (Chen and Vitetta 2020), increasing mucosal barrier integrity (Okumura et al. 2021), encouraging obligate anaerobic bacteria dominance (Chen and Vitetta 2020) and decreasing oxygen availability in the gut (Kelly et al. 2015). In an in vitro model of the intestinal epithelial barrier employing Caco-2 cells, SCFA, particularly butyrate, has been demonstrated to enhance intestinal barrier function as measured by an increase in transepithelial electrical resistance (TEER) and a decrease in inulin permeability (Peng et al. 2007), (Peng et al. 2009). Inflammatory bowel disease (IBD), obesity, non-alcoholic steatohepatitis (NASH), and non-alcoholic fatty liver disease (NAFLD) are all linked to a defective intestinal tight junction (TJ) barrier, which can be corrected by butyrate (Silva et al. 2018), (Coppola et al. 2021), (Endo et al. 2013). Reports showed that butyrate enhances the intestinal barrier by regulating the assembly of TJ proteins like occludin (Peng et al. 2009). Occludin, having four transmembrane domains is highly expressed at cell-cell contact sites and is important in the assembly and maintenance of TJ (Al-Sadi et al. 2011). Importantly, occludin knockout mice showed exhibited elevated inflammation, hyperplasia, and growth retardation (Saitou et al. 2000). Dysregulation of TJ has been implicated in various gut associated diseases including, IBD and colon cancer (Casalino et al. 2016). TJ protein expression at the cellular level is being governed by microRNAs, small regulatory RNAs (Ye et al. 2011). MiR122 is one of the microRNA that promotes Occludin mRNA decay by binding to its 3’UTR (Jingushi et al. 2017), (Yang et al. 2022). MiR122 is abundantly present in liver, but also reported to be found in gut where it targets NOD2 and plays important role in colon cancer (Li et al. 2019).

The study is designed to investigate the effect of prenatal arsenic exposure on gut functional phenotype in the post natal life. We studied the microbial composition, metabolite production and TJ protein expression and gut permeability in prenatally arsenic exposed mice and compared with normal control. Our results showed decrease in Firmicutes to Bacteriodes ratio in gut microbiota coupled with decrease in SCFA production of prenatally arsenic exposed mice. The lack of butyrate increased miR122 expression in gut causing decrease in occludin and resulted in increased gut permeability. By replenishing butyrate, the prominently decreased SCFA due to pre-natal arsenic exposed (pAs-mice) we showed that butyrate renders a crucial base in the maintenance of gut physiology. Further leverage on the role of miRNA122 was stemmed from over-expressing miR122 to butyrate treated prenatally arsenic exposed mice which led to decrease in occludin. By harmonising narratives from our experimental studies, we showed that in prenatal arsenic exposure, imbalance of the gut microbiota resulted in decrease in gut butyrate, a “critical” denominator for maintaining general gut homeostasis. We showed a lasting imprint of prenatal arsenic exposure on post-natal gut physiology in adult life even in absence of butyrate.

## 2. Materials and methods

### 2.1. Reagents, Chemicals, and Buffers

Chow diet (Harlan Teklad LM-485), was purchased from ICMR-NIN, Hyderabad, India. Dulbecco’s modified Eagle’s medium (DMEM) and foetal calf serum (FCS) were purchased from GIBCO (Waltham, MA, USA). BCA protein assay kit was purchased from Thermofisher (Waltham, MA, USA). Penicillin, streptomycin, Triton X100, PMSF, leupeptin, glycine, acrylamide, bis-acrylamide para-formaldehyde, sodium butyrate, sodium propionate, sodium acetate, Hoechst 33342, were purchased from Sigma (St. Louis, MO, USA). PVDF membrane, Trizol, were purchased from Invitrogen (Carlsbad, CA, USA). Super Reverse Transcriptase MuLV Kit, RT^2^ SYBR^®^ Green qPCR Mastermix were purchased from Qiagen (Hilden, Germany). Ripa lysis buffer, Occludin (rabbit, polyclonal) were purchased from Abcam (Cambridge, UK). Anti-GAPDH antibody (rabbit polyclonal) was purchased from Bio-Bharati (Kolkata, India). All primers were purchased from IDT (Lowa, USA). Human colon carcinoma cell line HT-29 cells were a kind gift from Dr. Amit Pal (ICMR-NICED, India). miR122 expression plasmid was a kind gift from Dr. Suvendranath Bhattacharya (CSIR-IICB).

### 2.2. Mice and Animal Ethics

Balb/c mice (6 weeks old) were procured from ICMR-NICED animal facility of the institute. All the protocol for the study was approved by the Institutional Animal Ethics Committee of ICMR-NICED, Kolkata, India, (PRO/151/July 2018–June2021). Experiments have been carried out in accordance with the guidelines laid down by the committee for the purpose of control and supervision of experiments on animals (CPCSEA), Ministry of Environment and Forests, Government of India, New Delhi, India.

### 2.3. Arsenic treatment to dams

All mice were housed in cages containing straw bedding held in pathogen-free facilities maintained at 24°C with a 50% relative humidity and 12-h light:dark cycle. All mice had ad libitum access to standard rodent chow. After 2 weeks of acclimatization, the Balb/c mice were bred by housing two females with a male and given ad libitum access to drinking water containing with/ without 4 ppm Arsenic trioxide (As) as described (Chakraborty and Bhaumik 2020). The water was changed twice weekly. After birth, the mothers were then given ad libitum access to clean As-free water. For the experiments, when pups reached 4week-of-age, groups were randomly collected, and processed for biomaterials. Age-matched juvenile mice whose mothers were never exposed to Arsenic were processed in parallel as controls. For each experiment, 5–6 juvenile mice were randomly chosen (without any sex bias) for evaluation.

### 2.4. Dietary supplementation of sodium butyrate

The dietary supplementation studies were performed as reported earlier with minor modification (Lin et al. 2012). Briefly, a group of 5 prenatally arsenic exposed mice pups (3 weeks old) were fed with 5% sodium butyrate (w/w) (Sodium butyrate in solid form were thoroughly blended into chow diet) for next 7 days (pAs-butyrate-mice) (Xu et al. 2018).

### 2.5. MiR122 overexpression in mice

MiR-122 was overexpressed in mouse gut in an identical procedure as described earlier (Ghosh et al. 2013). The miR-122 expression plasmid or empty plasmid (mock) was injected through the tail vein of pAs-butyrate-mice at a dose of 25 µg plasmid DNA dissolved in 100 µl saline. Following sacrifice of the mice after 4 days, the expression of occludin and miR122 in colon was determined by qPCR.

### 2.6. Determination of in vivo gut permeability

The gut permeability assay was performed following previously reported protocol (Rangan et al. 2019). Briefly, mice were starved 16 h prior to FD-4 administration. FD-4 (44 mg/100 g body weight) was administered by oral gavage with a needle attached to a 1 ml syringe. A gap of 30 min between each mouse was kept for the FD-4 oral gavage. After 4 h, the blood was collected from tail vein. The blood was immediately transferred to a Microtainer SST tube and was mixed by inverting the tube 3-4 times and was stored at 4 °C in the dark. Once blood was collected from all the mice, SST tubes are processed to separate the serum following the manufacturer’s instruction. The serum was then diluted with an equal amount of PBS. The concentration of FD-4 in sera were determined by spectrophotofluorometry with an excitation of 485 nm (20 nm band width) and an emission wavelength of 528 nm (20 nm band width) in Fluoroskan^TM^ Microplate Fluorometer (Thermo Fisher Scientific).

### 2.7. Mice fecal samples collection

Fresh fecal samples of all mice were collected at 15:00∼17:00 p.m. to minimize possible circadian effects. Samples were collected into empty sterile microtubes on ice and stored at −80 °C within 1h for future use.

### 2.8. 16S rRNA sequencing and analysis

For extraction of fecal DNA, fecal pellets were incubated for 24 hr at 56°C with proteinase K. DNA was then isolated using QIAamp DNA Mini Kits (Qiagen) using ∼25 mg of feces.

### 2.9. Evaluation of changes in the gut microbiome

Faecal pellets were collected from juvenile mice (irrespective of sex) from both the control and prenatally Arsenic exposed group. Bacterial genomic DNA was extracted using QIAamp Fast DNA Stool Mini Kit (Qiagen, Valencia, CA, United States) by following the manufacturer’s protocol. DNA concentration was evaluated by NanoDrop spectrophotometer from Thermo Fisher Scientific (Waltham, MA). Illumina standardized V3-V4 regions of 16s rRNA library protocol were employed for the preparation of the library. The library which was generated contained V3-V4 amplicons were then sequenced on an Illumina MiSeq platform following the manufacturer’s protocol. The generated reads (data) were processed by using QIIME2 (Bolyen et al. 2019) (version 2021.8.0). Filtering, merging and denoising was carried out by using DADA2 (Callahan et al. 2016) plugin within QIIME2. RESCRIPt (Robeson et al. 2021) plugin was used for processing and filtering of SILVA 138.1 (Quast et al. 2013) database to make it compatible with QIIME2 for carrying out taxonomy assignment. V3-V4 region primers were used for trimming SILVA sequencing. and classify-sklearn (Pedregosa et al 2011) was used for taxonomical classification. Further, the biome and taxonomy table was exported and diversity analyses were carried out by using phyloseq (McMurdie and Holmes 2013) in R Data. Alpha diversity and rarefaction curve were studied on samples (biom table) rarefy to a depth of 10,952 reads per sample. Samples generated during this experiment were submitted to Sequence Read Archives (SRA) of National Centre for Biotechnology Information (NCBI) under accession numbers (Control1=SRR19309430, Control 2= SRR19309429, pAs1= SRR19309428 and pAs2=SRR19309427). The raw data is available in bioproject_ accession PRJNA839617.

### 2.10. Estimation of faecal SCFA by GC/MS

The faecal concentrations of butyrate were measured by GC/MS as previously described (Li et al. 2020). Briefly, 50 mg of faecal samples from both groups were homogenized in 200 µl of distilled water. The samples were then centrifuges at 4000 rpm for 5 minutes and the resulting supernatant was collected. 200 µL of a benzyl alcohol–pyridine mixture (3:2) and 100 µL DMSO was added to the supernatant and the mixture was vortexed for 5 seconds. 100 µl of f benzyl chloroformate was added very carefully and the tube lids were kept open for 1 minute to release the gas formed during the reaction. The tube lids were then closed and the resulting mixture was vortexed for 3 minutes. After the derivatization, 200 µl of cyclohexane was added and the resulting mixture was vortexed for 1 minute which was then followed by centrifugation at 4000 rpm for 5 minutes. 100 µl of the resulting derivative extract (upper cyclohexane layer) was isolated and 1 µl was injected into GC-MS instrument for further analysis.

The samples were analysed using Shimadzu GCMS-QP2020 (Shimadzu Corporation, Kyoto, Japan) with an AOC-20i auto injector. An InertCap WAX capillary column (30 m × 0.32 mm × 0.25 μm; GL Sciences Inc., Tokyo, Japan) was used for separation. Helium was used as a carrier gas with a flow rate of 1ml/min. The temperature of the front inlet was set at 250 °C while the temperatures of the transfer line and the ion source were set to 280 °C and 230 °C respectively. The initial column temperature was held at 70 °C for 3 min and then was increased to 200 °C at a rate of 10 °C /min and was finally increased to 290 °C at a rate of 35 °C /min and then was held at this temperature for 7 min. A single run took 25.5 min. The solvent delay time was set to 6.7 min. The electron energy was −70 eV and the gain factor was set to 2.0.

### 2.11. Immunofluorescence Staining and imaging

Colon samples were collected from all the groups during necroscopy examination and were fixed in 4% paraformaldehyde. The sections were then embedded in paraffin and 5 μm sections were generated. The paraffin embedded sections were then de-paraffinized in xylene and were rehydrated by passing through a graded series of ethanol followed by rinsing with distilled water. Antigen retrieval was performed by immersing the slides in 1 mM EDTA buffer pH 8.0 for 5 minutes.at sub boiling temperatures. The slides were then washed with distilled water. The sections were then permeabilized by 0.1% sodium citrate and 0.5% Triton-X in TBST for 15 minutes. The sections were then blocked with 5% animal sera in TBST for 1 hour at room temperature. The primary antibody to Occludin was then diluted (1:200) in the blocking solution and added to the sections which were then incubated overnight in a humidified chamber at 4°C. Following incubation, the sections were washed TBS and TBST alternatively for 5 minutes. The sections were then incubated with goat anti-mouse or goat-anti rabbit secondary antibody conjugated with Alexa Fluor 488 for 2 hours at room temperature while being protected from light. Following incubation the slides were again washed with TBS and TBST alternatively for 5 minutes. The sections were then mounted with 1μg/ml Hoechst 33342 stain which acted as a nuclear counterstain. Fluorescence images were then captured using Carl Zeiss microscope equipped with a CCD camera and controlled by Zen software (Carl Zeiss, Gottingen, Germany) (Gumber et al. 2014).

### 2.12. Histopathologic examination of proximal colon

The proximal colon samples were fixed in 4% paraformaldehyde for 48 h at 4 °C. The fixed tissues were then dehydrated through graded alcohols, embedded in paraffin, and routine microtomy then carried out to generate 5 mM sections. The sections, in turn, were stained with hematoxylin and eosin for later microscopic examination. (Chakraborty and Bhaumik 2020)

### 2.13. Propagation of HT-29 cells and SCFA treatment

HT-29 cells were cultured in DMEM supplemented with 10% FCS and 50 µg/ml penicillin and streptomycin at 37° C with 5% CO_2_. Cellular viability was assayed by MTT. Cells (10^6^ cells/ml) were treated with either sodium butyrate (butyrate) or sodium propionate (propionate) or sodium aceatate (aceatate) at concentrations indicated in the figures for 24 h. Thereafter the cells were washed and processed for further analysis.

### 2.14. Tissue homogenisation and RNA and Protein isolation

Colon samples (dissected into small pieces) or the cells were resuspended either in RIPA Lysis buffer (20mM Tris-HCl pH 7.5, 150mM NaCl,, 1mM EDTA, 1mM EGTA, 1% NP-40, 1% Sodium deoxycholate, 2.5 mM Sodium Pyrophosphate, 1mM β-glycophosphate, 1mM Na_3_VO_4_, 1 µg/ml leupeptin with 1mM of PMSF immediately before use) for protein isolation or in Trizol (Invitrogen, US) for RNA isolation. The tissue was homogenized using a micropestle and centrifuged at 13,000 g for 15 min at 4°C. The clear supernatant was collected and either stored as protein lysate in −80°C or further processed to isolate RNA using the standard protocol (Wu et al. 2018). The concentration of the extracted RNA was analyzed by Nanodrop and RNA was stored at –80° C.

### 2.15. RNA extraction and reverse transcription

cDNA was prepared from total RNA by reverse specific primers using Super Reverse Transcriptase MuLV Kit. The primers for the reverse transcription are listed in Table 1. U6 and GAPDH were normalized for the expressions of miR122 and other genes of interest. The total reaction volume for reverse transcription was 20 μl in which 1 μM of reverse primer, 5 ng of RNA template, 1 μl dNTP mix, 12 μl of DEPC treated water, 4 μl of 5X first strand buffer, 1 μl of 0.1 M DTT, 1 μl of RNase inhibitor and 1 μl Super RT MuLV. Reverse transcription was carried out for 65°C for 5 minutes, followed by incubation at 55°C for 1 hour and then heat inactivating the reaction at 70°C for 15 minutes (Abdelmohsen et al. 2012).

### 2.16. Quantitative real-time PCR

Total RNA was extracted with Trizol reagent from snap frozen colon and RNA concentration was determined using a nanodrop. The miR-122, occludin, claudin1, claudin 2, claudin 4, ZO-1, GAPDH, and U6 levels were quantified with Applied Biosystems^TM^ StepOne^TM^ Real Time PCR System with RT^2^ SYBR^®^ Green qPCR Mastermix following the manufacturer’s instructions. Each 20 μl qPCR reaction contained an amount of cDNA equivalent to 5 ng of total RNA, 10 μl of RT^2^ SYBR^®^ Green qPCR Mastermix, 1 μM of the forward and reverse primer (each) and nuclease free water. Real-time PCR was performed with the following conditions: 95°C for 10 min, 40 cycles of 95°C for 30 sec, 60°C for 1 min and 72°C for 1 min PCR product was calculated according to the 2^^−ΔCt^ method described previously (Abdelmohsen et al. 2012).

### 2.17. Microbial analysis via PCR

To compare the relative abundances of various bacterial taxa in fecal DNA from female mice, we used qPCR analysis quantified by SYBR green and normalized the data to total bacterial abundance using pan-bacterial primers. The data is represented as 2^−Δct^ (Guo et al. 2008).

The primer sequences use for PCR amplification are as follows:

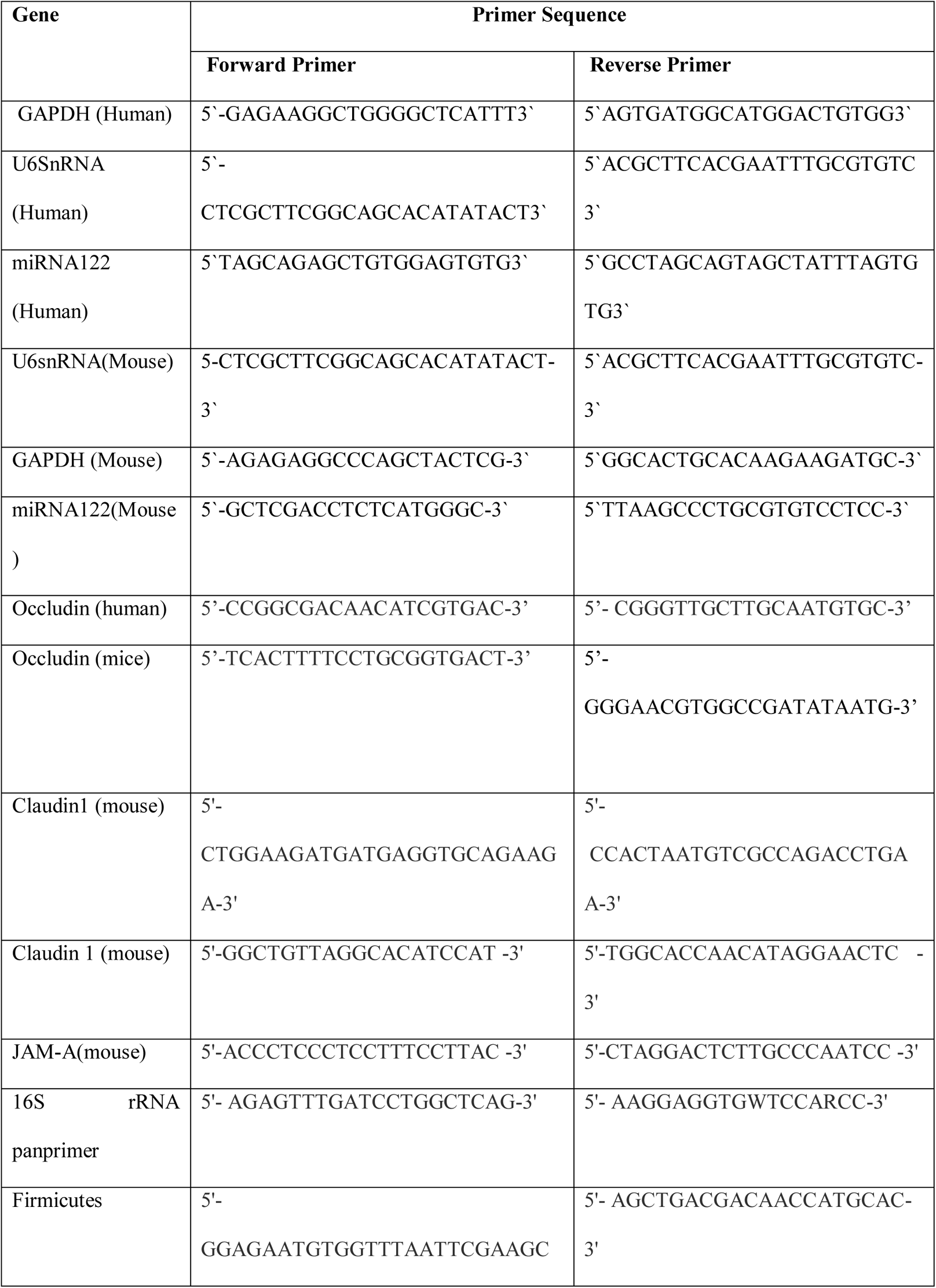
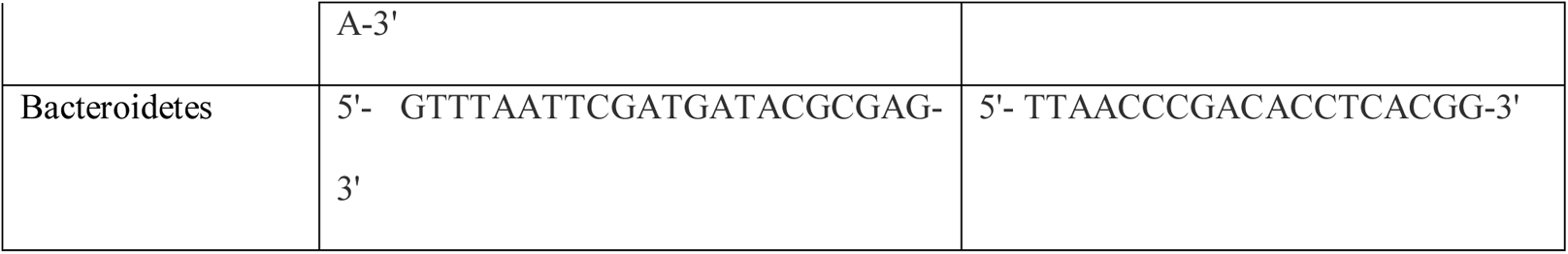

### 2.18. Western blot

Colon tissue protein and cell lysate were extracted in RIPA Lysis buffer. Protein concentration was measured using Pierce ^TM^ BCA Protein Assay Kit. Proteins (50 μg/lane) were separated by using SDS-PAGE on 10% gel under reducing condition and electro transferred to PVDF membrane in a transferred buffer (25mM Tris-HCl, 150mM Glycine, 20% Methanol). Membranes were blocked at room temperature with 5% non fat skim milk in TBS for 2 hours, and then incubated with primary antibody against specific protein. The membranes were incubated with the horseradish peroxidase (HRP)-conjugated secondary antibodies at 37° C for 1 h. SuperSignal West Pico chemiluminescent substrate kit (Thermo) was used to visualize the blotting results. The blots were imaged with Fluor Chem R system (ProteinSimple, San Jose, CA, USA) (Abdelmohsen et al. 2012).

Antibody used for Western Blots

2.18. Western blot Antibody used for Western Blots
| Name of Antigen | Raised in | Source | Dilutions used |
| --- | --- | --- | --- |
| Anti-mouse-HRP<br>conjugated secondary<br>antibody | Horse | Cell Signalling Technology | 1:5000 |
| Anti-rabbit- HRP<br>conjugated secondary<br>antibody | Goat | Cell Signalling Technology | 1:5000 |
| Occludin | Rabbit | Abcam | 1:1000 |
| GAPDH | Rabbit | Biobharati | 1:1000 |

### Statistical Analysis

All values in the figures and text are expressed as arithmetic mean ± SEM. Data were analyzed with GraphPad Prism Version 8.01 software and statistical significance between groups was determined using unpaired student’s t-test. The *p* values of *<*0.05 were considered statistically significant. In the experiment involving Western blot, the figures shown are representative of at least 3 experiments performed on different days.

## 3. Results

### 3.1. Arsenic trioxide (arsenic) in drinking water decreases Firmicutes and increases Bacteroidetes in female mice

Female Balb/c mice were fed with 4 ppm arsenic trioxide (arsenic) in drinking water since breeding and continued till parturition (As-exposed-dams). Another group of female mice fed with normal drinking water (without arsenic) were set for breeding and served as control group (Control-dams). Both the groups were fed with food and water *ad libitum*. We investigated whether arsenic would perturb the abundance of *Firmicutes* (F) and *Bacteroidetes* (B) present in the gut microbiome of the dams. The abundance of *Firmicutes* and *Bacteroidetes* was measured by qPCR using phyla specific primers. The relative abundance was calculated by normalizing with bacterial pan specific primer. We observed 2.5 fold decrease (p=0.04) in *Firmicutes* and 7 fold increase (p=0.0008) in *Bacteroidetes* in As-exposed dams (Fig S1 A and B) resulting to 18 fold decrease (p=0.005) in F/B ratio compared to control (Fig S1).

**Figure 1:**
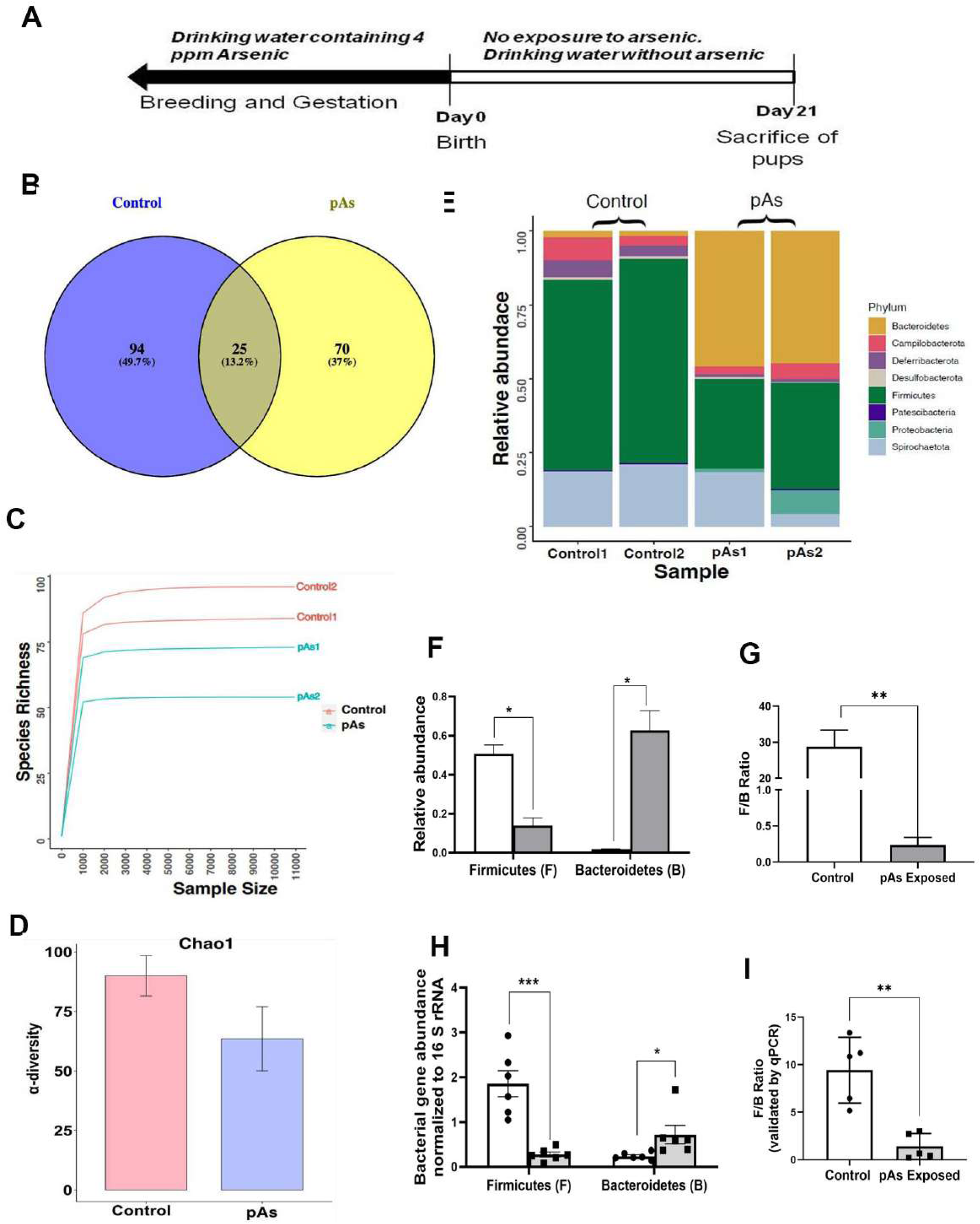
Effect of prenatal Arsenic exposure on gut microbiome composition and diversity. Adult Balb/c mice were bred by housing two females with a male and given ad libitum access to drinking water containing 4 ppm arsenic trioxide (As). After birth of the pups, the mothers were then given ad libitum access to clean As-free water. When the pups reached 4weeks-of-age, their feaces were collected at random. Feaces from the age-matched pups of the dams that were never exposed to Arsenic were processed in parallel as control. The experimental plan is schematically represented (A). Venn-Diagram showing the number of microbial families’ common and exclusive to control (control-mice) and prenatally As exposed mice (pAs-mice) (B). Species rarefaction curves outlining the number of bacterial species in control and pAs mice samples. (C). The Chao1 α-diversity of microbial communities in control and pAs mice (D). Stacked histograms are showing major phylum abundant in control and pAs mice. (E). Relative Abundance of Firmicutes and Bacteroidetes in control and pAs exposed mice (F). Firmicutes (F) to Bacteroidetes (B) ratio in control and pAs exposed mice (G). Firmicutes and Bacteroidetes abundance as measured by qPCR (H) and F/B ratio (I) (n=6/group for qPCR and n=2/ group for 16S rRNA sequencing). Data represented as mean ± SE. Values significantly different from control at *p<0.05, **p<0.01 and ***p<0.001.

Next we asked whether arsenic exposure during breeding and gestation would perturb the compositional profile of the gut microbiome of the offspring.

### 3.2. Prenatal arsenic exposure perturbs normal community composition of gut microbes

The pups from the As-exposed-dams were not exposed to arsenic any further after their birth and referred to as pAs-mice henceforth for convenience. The pups from the control-dams were referred as control-mice. The feaces were collected from the pups on day 21 post birth as represented pictorially (Fig 1A). The total number of taxonomically classified ASVs (Amplicon Sequence Variants) identified upon analysis of sequenced 16S-rRNA dataset from the feaces in both the groups is represented in a Venn diagram (Fig 1B). The identified genera are enlisted in Table S1. Around 13.2% ASV was found to be common in both the groups. But control-mice and pAs-mice showed the presence of 49.7 % and 37% unique ASV respectively. Overall this observation indicates pAs-mice have lost about 12.7% of ASV compared to control-mice (Fig 1B). The predominant bacterial phyla in all the samples were *Firmicutes* and *Bacteroidetes* as assessed by 16S-rRNA sequencing (Fig 1E). Alpha diversity of the samples was measured by chao1 index. Following prenatal arsenic exposure, the microbial component of the gut microbiome was lowered significantly compared to control, along with a Species Rarefaction Curve (Fig 1C) showing reduction in the number of bacterial species in the pAs mice (line in red) with respect to control-mice (line in blue) and a decrease in diversity (chao1 value for sample: Control1=84, Control2=96, pAs1=73, pAs2=54) (Fig 1D). At the phylum level, the prevalence of *Firmicutes* and *Bacteroidetes* as determined by metagenomic analysis were significantly decreased and increased respectively in the pAs-mice compared to control-mice (Fig 1F). Considering the decrease in *Firmicutes* (F) and increase in *Bacteroidetes* (B) in pAs-mice, the F/B ratio is also decreased (p=0.0254) in pAs-mice compared to control (Figure 1G). The abundance of these phyla were further validated by qPCR using phyla specific primers which also showed decrease and increase in *Firmicutes* and *Bacteroidetes* respectively in pAs-mice compared to control-mice (Fig 1H). We report 6.67 fold decreases (p=0.0013) in F/B ratio in pAs-mice compared to control (Fig 1I).

### 3.3. pAs-mice showed reduced SCFA production compared to control-mice

So far from the results it was apparent that there was a reduction in SCFA producing bacteria in the gut of pAs-mice compared to control. Therefore, we estimated the fecal SCFA content of control and pAs mice. The decrease in the abundance in *Firmicutes* in pAs mice faithfully reflected decrease in fecal SCFA compared to control as estimated by GC-MS. All the SCFAs including propionate, aceatate and butyrate were decreased significantly compared to pAs-mice. We observed 4 fold (p=0.0079), 2 fold (p=0.0049) and1.7 fold (p=0.025) decrease in butyrate, aceatate and propionate respectively in the feaces of pAs-mice compared to control-mice (Fig 2).

**Figure 2:**
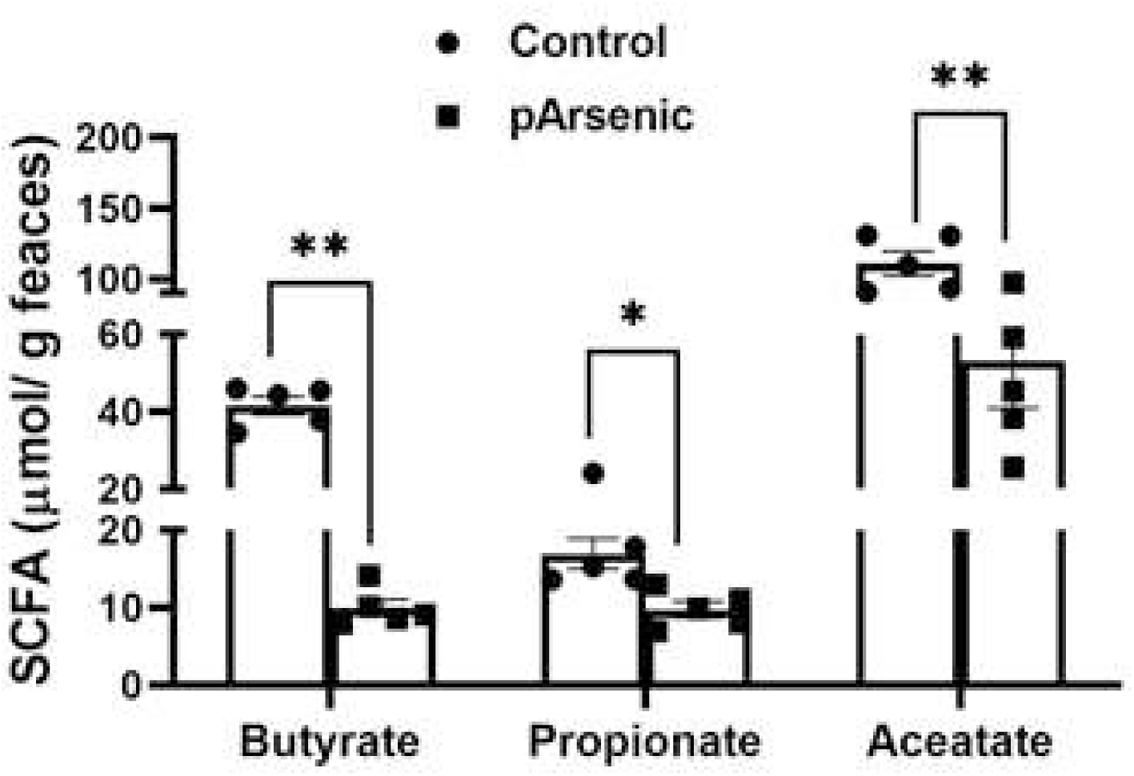
Effect of prenatal arsenic exposure on concentration of faecal short chain fatty acids. Amount of faecal butyrate, acetate and propionate in control-mice and pAs-mice was determined by GC-MS. Data represented as mean ± SE (n=5/group) *p<0.05, **p<0.01 and ***p<0.001.

As SCFAs perform multiple functions including intestinal barrier integrity, mucous production and protect against inflammation in gut it was important to study the physiological function of the gut in pAs-mice.

### 3.4. pAs-exposed mice showed increase in gut permeability and decrease in tight junction (TJ) protein, Occludin

To assess the potential shifts in the gut physiological function, if any, the gut barrier function, junctional protein expression and histology of gut in pAs-mice and control-mice were studied. The gut barrier function as studied by FITC dextran permeability showed 3.9 fold increase (p=0.0079) in gut permeability in pAs-mice compared to the control-mice (Figure 3A). Further evaluation of tight junctional protein expression in the colon by qPCR showed 4 fold decrease (p=0.007) in Occludin in pAs-mice compared to control-mice whereas, claudin 1, claudin-2, claudin-4, ZO-1 and JAM-A TJ protein expression remained unaltered (Fig 3 B). Histological examination of the gut showed inflammatory cell infiltration in pAs-mice compared to the control-mice (Fig 3C). The mucous secreting goblet cells also showed hyperplasia in pAs-exposed animals’ gut histological sections.

**Figure 3:**
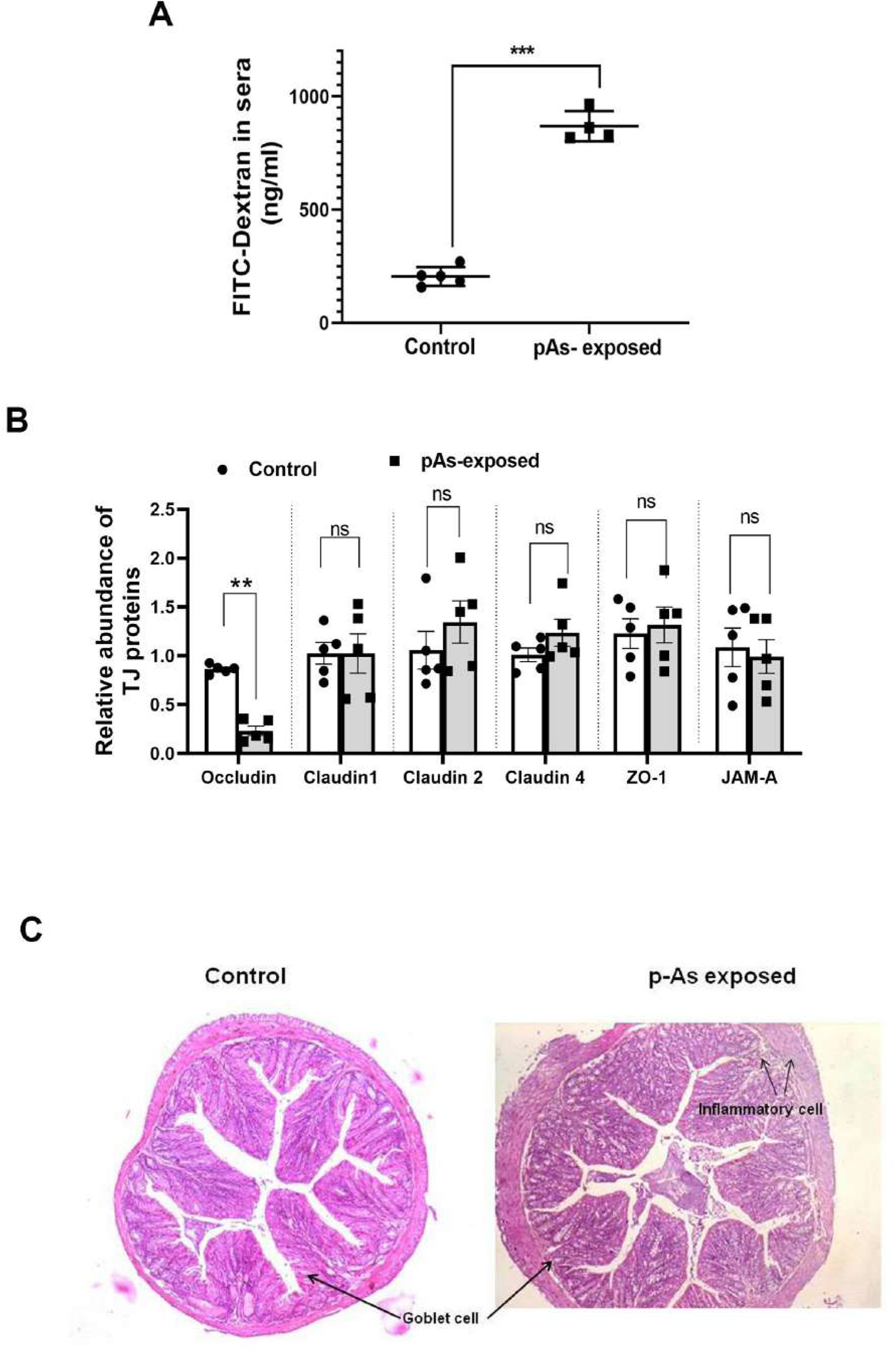
Effect of prenatal Arsenic exposure on intestinal histoarchitecture and intestinal permeability. Control-mice and pAs-mice were starved for 16 h and then fed with FITC-Dextran 4000 (FD4) by oral gavage at a dose of 44 mg/100 g body weight. After 4 h the blood was collected and serum was extracted to determine the fluorescence of FD-4 in sera (A). The expression of tight junction proteins (Occludin, Claudin-1, Claudin-2, Claudin-4, ZO-1 and JAM-A) as determined by qPCR from colon tissue of Control-mice and pAs-mice (B). Representative micrograph of colon histology of control and pAs-mice (H&E; 10x magnification) (C). Arrows showing neutrophil infiltration. Data represented as mean ± SE. N=5/ group. Values significantly different from control at **p< 0.01 and ***p<0.001.

Recalling the gut microbial analysis and production of SCFA which depicted reduced SCFA production in pAs-mice, we sought to study if the lack of SCFA production in gut has any role on occludin expression in pAs-mice. Before that it was essential to investigate the mechanism by which SCFA may influence the expression of occludin (if any).

### 3.4. Butyrate treatment increases Occludin expression and decreases permeability in HT-29 cells

HT-29 cells were treated with either butyrate or propionate or aceatate with the function of concentration (0-20 mM) for 24 h. It was observed that unlike propionate and acetate (Fig S3), 10mM and 20mM butyrate treatment caused 45% and 58 % increase in occludin expression respectively (Fig S2). These results provided initial evidence that butyrate but not propionate or aceatate could increase occludin expression. For further deciphering the mechanism of regulation of occludin expression we studied miR122 expression upon butyrate treatment as it was shown previously that miR122 binds to the 3’UTR of the mRNA of Occludin, causing its degradation (Jingushi et al. 2017).

### 3.5. Butyrate treatment reduces miR122 expression in HT-29 cells

We monitored the status of miR122 in butyrate treated and untreated HT-29 cells by qPCR. It was observed that there was a progressive decrease in miR122 expression as a function of butyrate concentration (Fig S2). Notably, the expression of miR122 does not change on propionate and aceatate treatment. Our result points towards the cascade of sine-qua-non events accomplished by butyrate action which is as follows: butyrate-miR122-occludin which was further evaluated in pAs-exposed mice.

### 3.6. Butyrate treatment recovers gut permeability by reducing miR122 and increasing Occludin expression in pAs-exposed mice

To show that decrease in gut-derived butyrate is intimately linked with decrease in barrier function in pAs mice, we supplemented butyrate orally to pAs mice and studied the reversal of barrier function (if any). With this objective, we fed the pAs-mice from day 21-28 with 5% butyrate mixed with diet. The expression Occludin in the gut was studied by western blot in control-mice, pAs-mice and pAs-exposed mice fed with butyrate (pAs-butyrate-mice) (Fig 4A). The results showed that there was significant down regulation of occludin expression in the gut of pAs-mice which returned to normal with butyrate treatment (Fig 4B). Similarly the immunohistochemical analysis showed that occludin protein expression in the gut samples of pAs-mice was 2.8 fold reduced compared to control-mice which were restored to normal in pAs-butyrate-mice (Fig 4C, D). The miR122 expression showed 3 fold increases in pAs-mice compared to control and after butyrate treatment the miR122 expression essentially decreased compared to pAs-mice and was equivalent to control (Fig 4E). As expected, there was significant increase in permeability in pAs-mice compared to control. Although in pAs-butyrate-mice the gut permeability decreased significantly compared to pAs-mice, it remains higher in respect to control-mice (Fig 4F).

**Figure 4:**
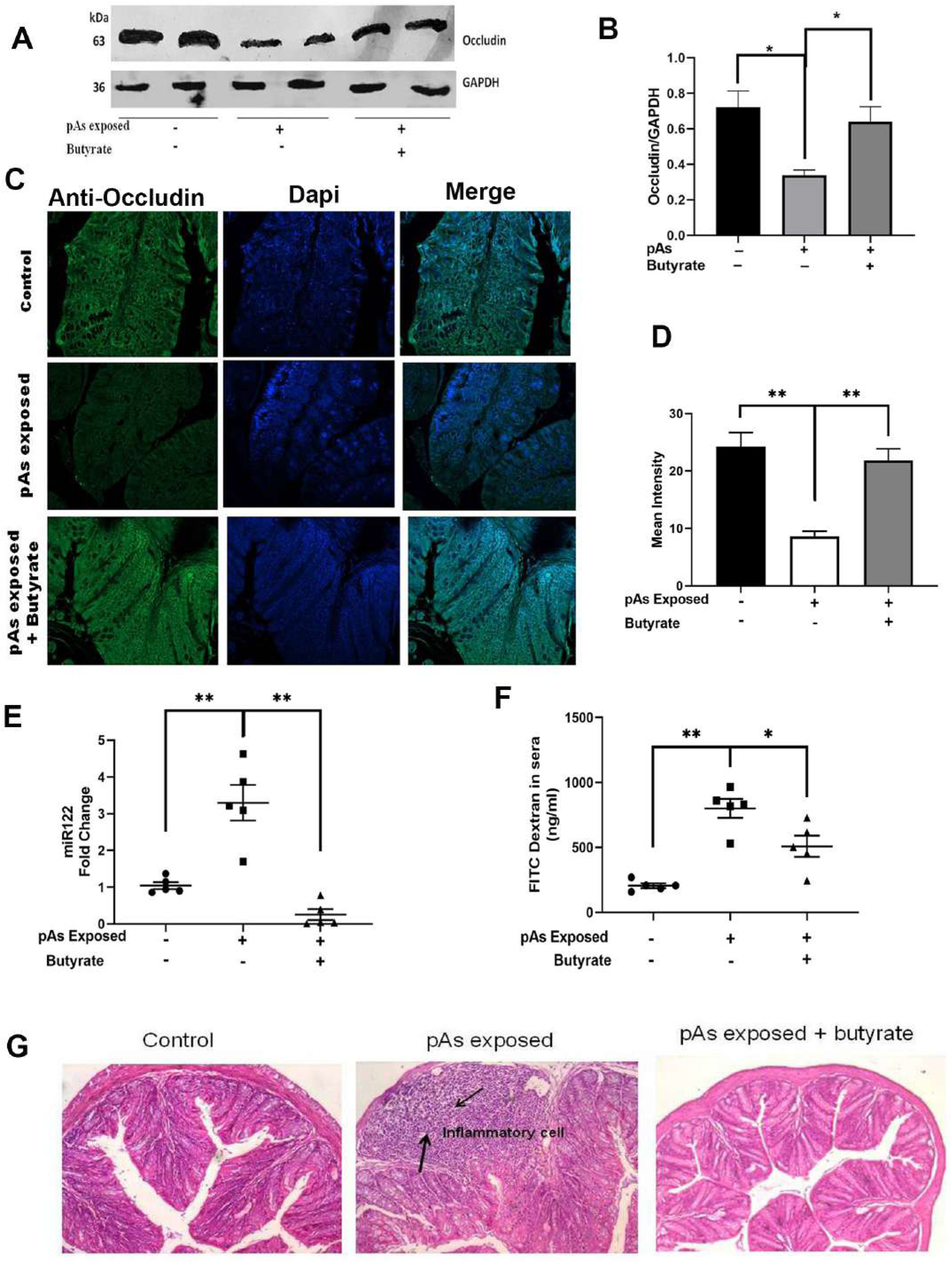
Effect of oral supplementation of butyrate on colon histoarchitecture and intestinal permeability of prenatally arsenic exposed mice. Expression of Occludin as studied by Western blot **(A)** showing corresponding densitometry (B) and Immunofluorescence (C) showing mean intensity (D), in control-mice, pAs-mice and pAs-butyrate-mice. Expression of miR-122 quantified by qPCR in control-mice, pAs-mice and pAs-butyrate-mice (E). Intestinal permeability as measured by the presence of FITC-Dextran 4000 (FD4) in serum in control-mice, pAs-mice and pAs-butyrate-mice (F). Representative micrographs of colon sections from control-mice, pAs –mice and pAs-butyrate –mice (H&E; 20x magnification) (G). Arrows indicating neutrophil infiltration. Data represented as mean ± SE. N=5/ group. Values significantly different from control at *p<0.05 and **p<0.01.

The colon histology sections revealed that butyrate restores gut heath by inhibiting recruitment of inflammatory cells like neutrophils that was induced by pAs-exposure (Fig 4G). Furthermore infiltration of inflammatory cells and goblet cell hyperplasia were reduced in pAs-butyrate-mice.

### 3.7. MiR122 over-expression rescues the effect of butyrate in pAs-butyrate-mice

From the previous experiments it was indicated that butyrate downregulates miR122 and upregulates occludin expression in colon. Hence, it was essential to confirm our observation by over-expressing miR122 in pAs-butyrate-mice. The miR122 expressing plasmid was injected through tail vein (25 µg/mouse) to the pAs-butyrate-mice. Another group of mice were injected with mock plasmid and day-4 post injection the animals were sacrificed and the gut was collected for further analysis. The level of miR122 and occludin expression in the colon of control-mice, pAs-mice and pAs-butyrate-mice were similar to the previous experiment. MiR122 over-expression in pAs-butyrate-mice showed high levels of miR122 in colon indicating successful overexpression of miR122 (Fig 5A). Corresponding analysis of occludin expression in the colon of pAs-butyrate-mice showed significant decrease with miR122-plasmid injection compared to mock-plasmid injection as studied by qPCR and western blot (Fig 5B, C, D).

**Figure 5:**
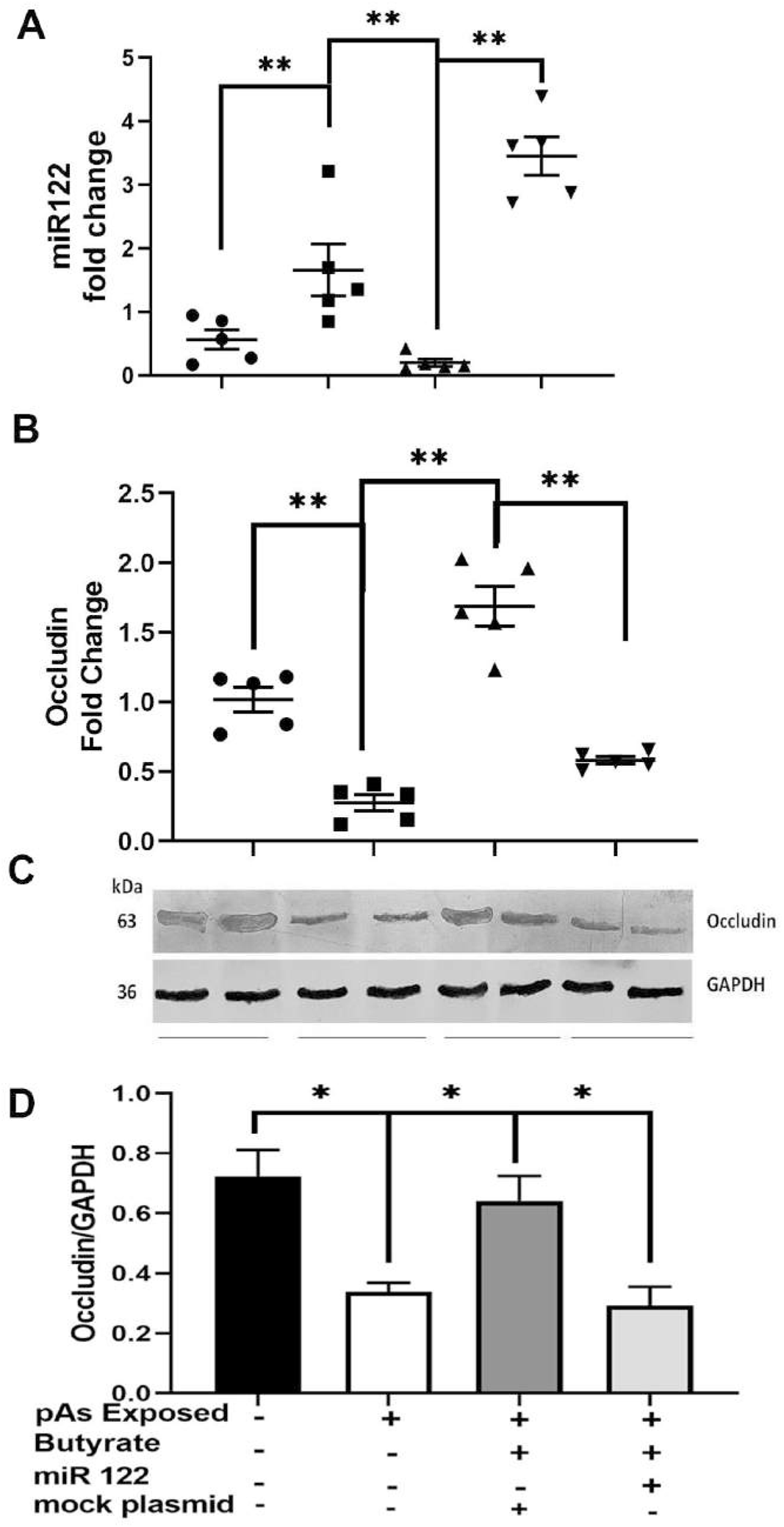
Effect of overexpression of miR-122 on Occludin expression in mice colon. Each butyrate treated pAs-mice were injected with either 25 µg in 100 µl of miR122 expressing plasmid or 25 µg in 100 µl of mock plasmid in tail vein. The mice were sacrificed 4 days post injection. The miR122 (A) and occludin (B) expression in colon was measured. N=5, the data is represented as mean ± SE. The experiment was repeated twice. **** represents p<0.0001, *** represents p<0.001, ** represents p<0.01.

## 4. Discussion

The ambit of this investigation lies in answering an impending question whether *in utero* arsenic exposure can affect the gut microbiome which may have caused altered gut physiology. Earlier reports showed that the environmentally relevant level of arsenic perturbs the normal gut microbiome in mice, decreasing abundance of *Firmicutes*, increase in *Bacteroidetes* and also changing its metabolites particularly SCFA (Chi et al. 2017). We validated this observation and showed that oral arsenic treatment causes decrease in *Firmicutes* and increase in *Bacteroidetes* in the dams. Considering the possibility of mother-offspring transmission of gut microbiome (Van Daele et al. 2019) we compared the broad microbiome composition of pAs-mice and control-mice. Overall, around 12.7 % ASVs (Amplicon Sequence Variant) of normal gut bacteria was found to be lost in pAs-mice compared to control-mice. Arsenic is shown to prolong glycan residues of cell membrane glycoprotein of skin cancer cells (Lee et al. 2016). Glycosylation of the intestinal mucus and epithelium is quite complex and can change in response to microbial colonization (Pickard et al. 2017). This is interesting because host glycans can serve as nutrient sources or adhesion receptors for microbes (Pickard et al. 2017). Therefore it is tempting to speculate arsenic may reshape gut microbial composition in the mice by changing the extent of glycosylation pattern of the receptor proteins in the epithelial cell membrane. Diversity of a community depends on the intensity of sampling and Chao1 is measured based on species richness (number of species in the community) in the groups. Although not significant, Chao1 alpha diversity of the microbiome showed a trend of decrease in pAs-mice compared to control-mice. In some cases, microbial compositions rather than diversity seems to be the key player to determine the phenotype as it was shown earlier with the survivality of wild vertebrate population (Worsley et al. 2021).

Our further analysis of the phylum composition of gut microbiota in pAs-mice and control-mice showed a significant shift in the relative abundance of *Firmicutes* (F) and *Bacteriodetes* (B) by metagenomics analysis and also by qPCR. Surprisingly, the extent of decrease and increase of *Firmicutes* (F) and *Bacteriodetes* (B) respectively in pAs-mice compared to control-mice as determined by the two methods were not hand in hand. This discrepancy may be due to low sensitivity of qPCR compared to metagenomics (Plaire et al. 2017) (Andersen et al. 2017). We report decrease in *Firmicutes* (F) and increase in *Bacteriodetes* (B) leading to consolidated decrease in F/B ratio in the gut communities of pAs-mice compared to control-mice which essentially mirrors the F/B ratio of the dams. The difference in the magnitude of the drop in F/B ratio between dams and offsprings could be attributable to the dams’ vaginal microbiome and environmental factors that also influence the establishment of gut microbiome in neonates (Tudor et al. 2017). The decrease in F/B ratio is a major biomarker of Inflammatory Bowel Disease (IBD) (Guo et al. 2021) suggesting a possibility that pAs-mice may have physiological dysfunction in the gut. The decrease in the abundance of *Firmicutes* in pAs-mice foretell the decrease in SCFA (den Besten et al. 2013). As expected, we observed significant decrease in all SCFAs including aceatate, propionate and butyrate in the feaces of pAs-mice compared to control-mice. Notably, the decrease in butyrate was more pronounced than the decrease in aceatate and propionate. The perturbations in the bacterial community and metabolites has overarching effect on the gut physiological functions as SCFAs particularly butyrate plays important role in multiple physiological processes in the host (Canani et al. 2011). Butyrate is an important energy source for intestinal epithelial cells (Singh et al. 1997) that maintains colonic homeostasis (Gasaly et al. 2021) and inhibits inflammation (Segain et al. 2000).

The gastrointestinal epithelium forms the body’s largest interface with the external environment (Groschwitz and Hogan 2009). It effectively provides a barrier that selectively limits permeation of luminal toxins and antigens through the mucosa (Suzuki 2013). The physical location of the intestinal epithelium, which is wedged between the luminal contents and the mucosal surface, supports the notion that a breach in the mucosal barrier causes mucosal inflammation (Ahmad et al. 2017). Studies with knockout mice with TJ proteins, develop inflammation in the gut epithelium (Ahmad et al. 2017), (Lu et al. 2013) further studies support the key role of permeability, especially in its capacity to contribute to overall mucosal barrier function in regulating mucosal immune homeostasis (Castoldi et al. 2015).

Pertinent with this idea we studied the corpus of events of gut physiological function which revealed increased permeability in gut and down regulation of tight junction protein, Occludin whereas other TJ proteins like Claudin 1, claudin-2, claudin 4, ZO-1 and JaM-A remain unaltered. Tight junctions are complex signalling centres in a continually changing milieu (Weber 2012) serving as a permeability barrier, preventing free passage of solutes via the intercellular space. Claudins, in combination with the cytoplasmic scaffold ZO, create TJ strands and perform critical roles in epithelial barrier assembly (Furuse 2010). In addition to claudins, TJs are home to other integral membrane proteins such as occludin, a tetraspanning membrane protein, and immunoglobulin superfamily proteins, including junctional adhesion molecules (JAMs) (Furuse 2010). They also play an important role in the regulation of paracellular permeability. The coiled coil domain of occludin acts to organize the structural and functional elements of TJ (Nusrat et al. 2000). Paracellular permeability is, to a great extent, controlled by tight junctions, and disrupting their integrity, assembly and expression results in increased permeability (Liang and Weber 2014). Occludin being an important component of tight junction, the decrease in its expression or function leads to increase in permeability. The translocation of luminal components into the host could cause both local and systemic inflammatory pathways in the case of enhanced intestinal permeability (Mu et al. 2017). Arsenic treated Caco2 cells were shown to increase paracellular permeability by redistribution of zona occludens and reduced claudin 1 expression (Chiocchetti et al. 2019). In contrast, our study showed that prenatal arsenic exposure does not change the expression of claudin 1 and ZO-1. Possibly, as the pAs-mice were not exposed to arsenic after birth the toxic concentration of arsenic that is required to reduce claudin1 is not attained in the juvenile body.

Recent report showed arsenic treatment impairs distinct population of intestinal stromal cells and intraepithelial and innate immune cells (Kellett et al. 2022) (Medina et al. 2020). In conjunction, the histology of colon sections of pAs-mice showed infiltration of inflammatory cells and goblet cell hyperplasia. Gut barrier disruption and neutrophil infiltration are closely associated phenomena (Lin et al. 2020). The precise mechanism of goblet cell hyperplasia is unclear. A previous study showed that IL-13, the key regulator in type-2 mediated inflammation induces goblet cell hyperplasia to accelerate inflammation (Huang et al. 2020). Arsenic has been linked to IL-13 induction (Rahman et al. 2021), which could possibly lead to goblet cell hyperplasia.

SCFA, particularly butyrate has been shown to strengthen barrier function and decrease intestinal permeability in several studies using cell culture models and animal models (Peng et al. 2009). Recalling the decrease of SCFA producing bacteria and SCFA production in pAs mice we studied if SCFA has any effect on occludin expression. For deeper understanding of the effect of SCFA on occludin expression we undertook studies in colon carcinoma cell line, HT-29. The concentration of SCFA used in the treatment of HT-29 cells is effectively luminal concentration (Liu et al. 2018). Butyrate but not propionate or aceatate treatment to HT-29 cells showed increase in Occludin expression. To further understand the mechanism by which butyrate regulates occludin expression we sought to study miRNA that plays a crucial role in regulating gene expression (Kaikkonen et al. 2011). It is reported that miR122, binds to the 3’UTR of Occludin mRNA causing its decay (Jingushi et al. 2017). Although miR122 abundantly found in liver, it is also reported to express in the intestinal tissue (Runtsch et al. 2014). Our study showed that butyrate decreases miR122 expression as function of its concentration. To confirm the inhibition of miR122 is specific to butyrate and not other SCFAs we showed that neither propionate nor aceatate changes miR122 and Occludin expression. In another study from our group we have shown that a RNA binding protein, AUF1 plays a fulcrum point in miR122 regulation by butyrate (data not shown and the manuscript is under review).

Our findings capture transitivity of miR122 and Occludin in butyrate mediated decrease in paracellular permeability. The elegance of the interaction of these molecular players was further verified by providing butyrate orally in pAs-mice. As expected, butyrate treatment to pAs-mice, Occludin expression in gut was increased with concomitant decrease in gut permeability. Butyrate fed pAs-mice not only restored miR122 level and occludin expression which resulted decreased gut permeability and reduced infiltration of neutrophil in the gut. Further to understand the molecular events associated with butyrate and Occludin expression we over-expressed miR122 in butyrate treated pAs-mice resulted in appreciable recovery of Occludin expression.

The limitation of our study is that it is focussed only at a particular age of the post natal life. Further longitudinal studies are required to understand how microbial composition and gut functions changes with age in the post natal life if not exposed to arsenic anymore. Using technology involving CRISPR-based recording method by *E coli* sentinel cells to reveal transcriptional changes in intestinal and microbial physiology (Schmidt et al. 2022) will provide additional perspectives of arsenic induced changes in future.

## 5. Conclusion

Overall, the present study deals with an interesting connection of prenatal arsenic exposure and altered gut physiology in post natal life. We report gut microbial dysbiosis in pAs-mice leading to decrease in Firmicutes to Bacteroidetes ratio and decrease in production of SCFAs. We also document increase in gut permeability with decrease in Occludin expression which was reversed after butyrate treatment. Breach in the gut barrier function increases inflammatory gene expression in the gut of pAs-mice which was further reversed by butyrate treatment. Employing in vitro and in vivo experiments we have shown that butyrate down regulates miR122 expression which is responsible for increase in Occludin expression leading to the decrease in permeability. By rescuing miR122 expression after butyrate treatment we further establish the sequential molecular partners-miR122 and Occludin that plays a role in butyrate mediated increase in barrier function in prenatal arsenic exposed mice.

## Acknowledgement

We acknowledge the support of Director, ICMR-NICED, Kolkata for carrying out the study. We thank Dr. P. Jaisankar, Sudip Dey and Sandip Chaudhury (CSIR-IICB) for GC-MS analysis. We also thank Dr. Anjan Das for analyzing the histology. We acknowledge Prof. Syamal Roy (CSIR-IICB) for critically reviewing the manuscript. MC and OD are recipient of fellowship from the CSIR and DST respectively.

## SUPPLEMENTARY

**Figure S1:**
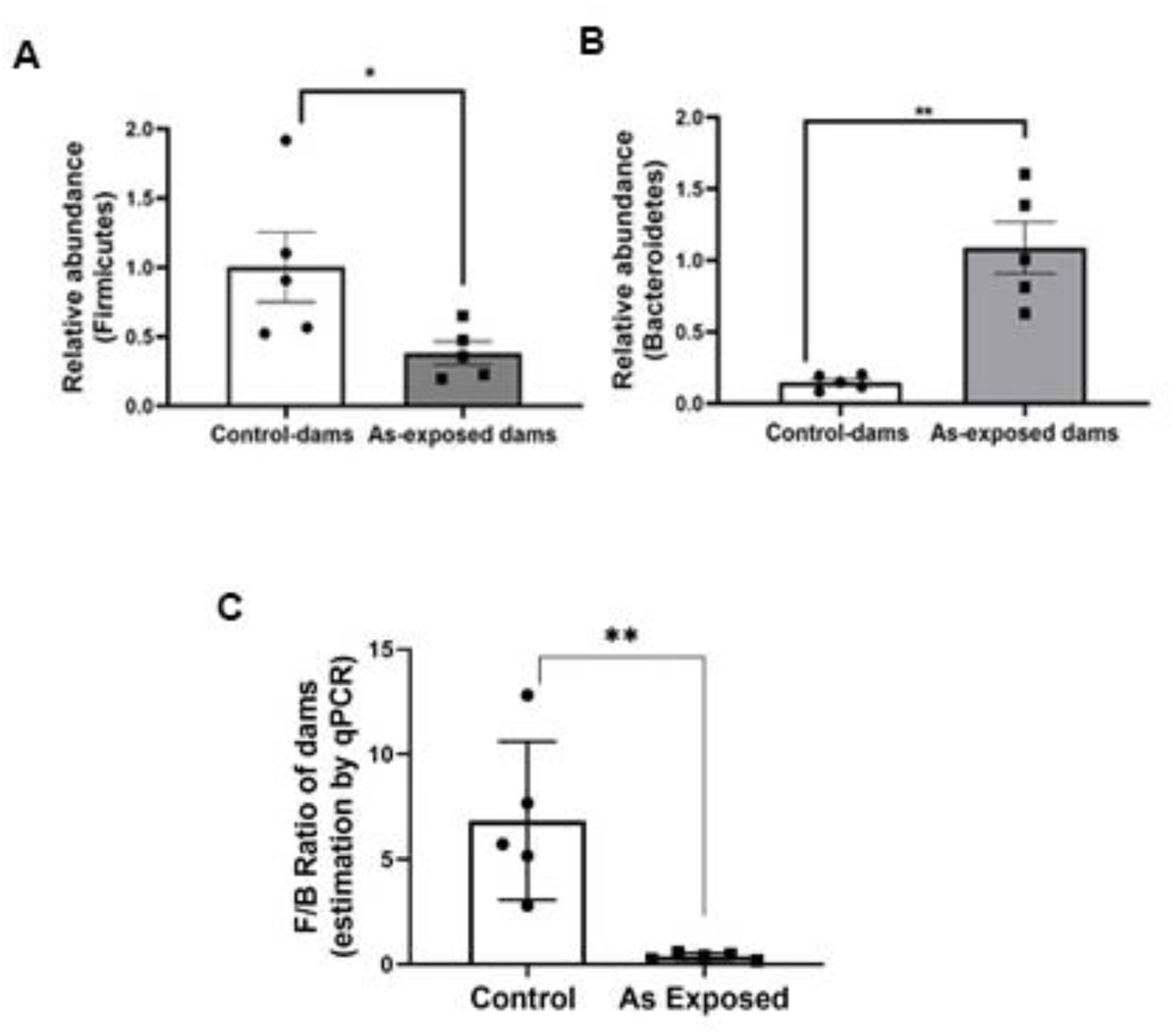
Adult balb/c mice were bred by housing two females with a male and given ad libitum access to drinking water containing 4 ppm arsenic trioxide (As). The feaces were collected from the pregnant dams 2-3 days before the birth of the pups. The relative abundance of Firmicutes (A) and Bacteroidetes (B) and F/B ratio (C) from control-dam and As-dams were measured by qPCR.

**Figure S2:**
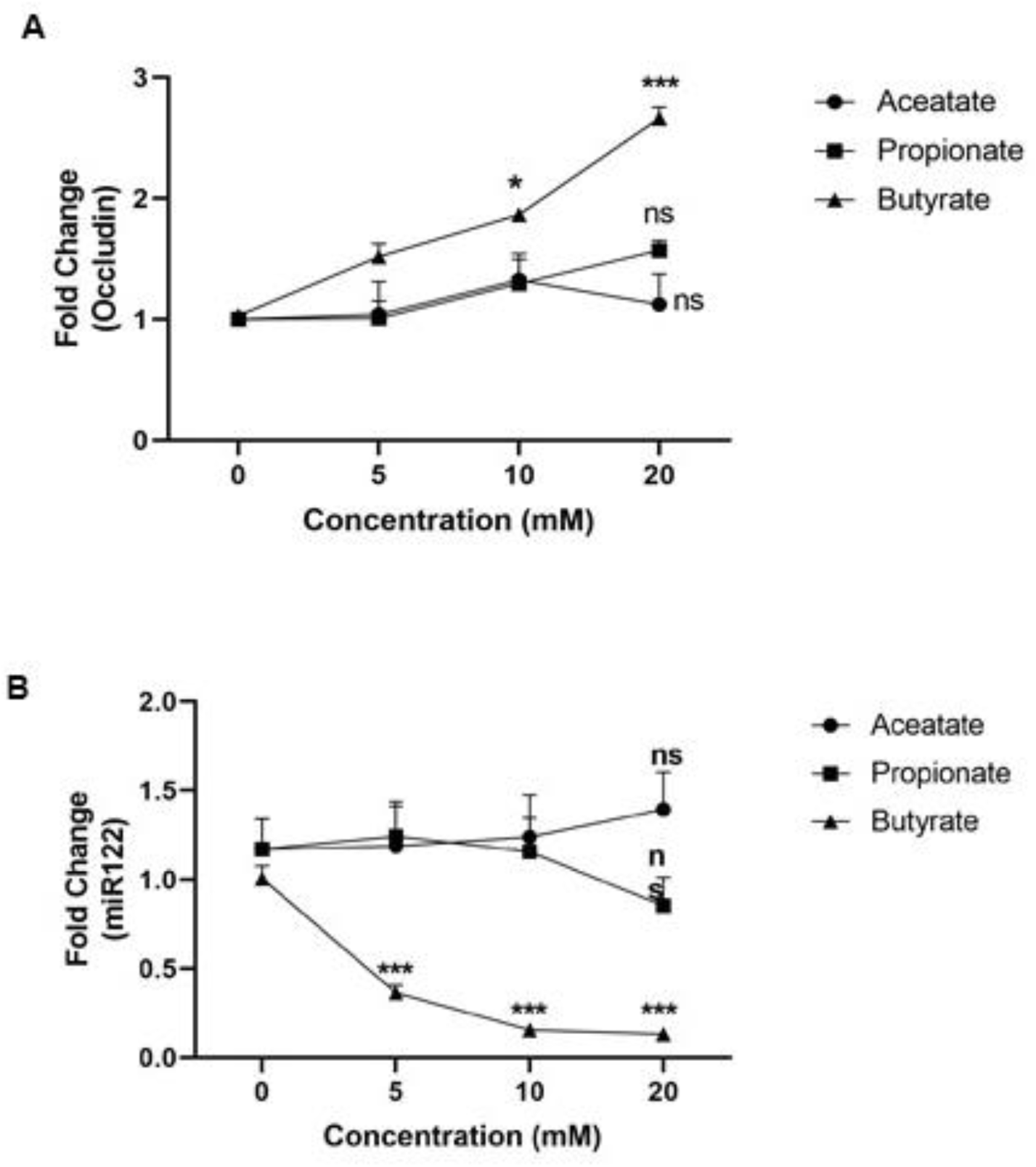
Expression of Occludin (A) and miR-122 (B) in HT29 cells treated with butyrate, propionate and acetate at varying concentrations (5mM, 10mM and 20mM) for 24 h as measured by qPCR.

**Table S1:**
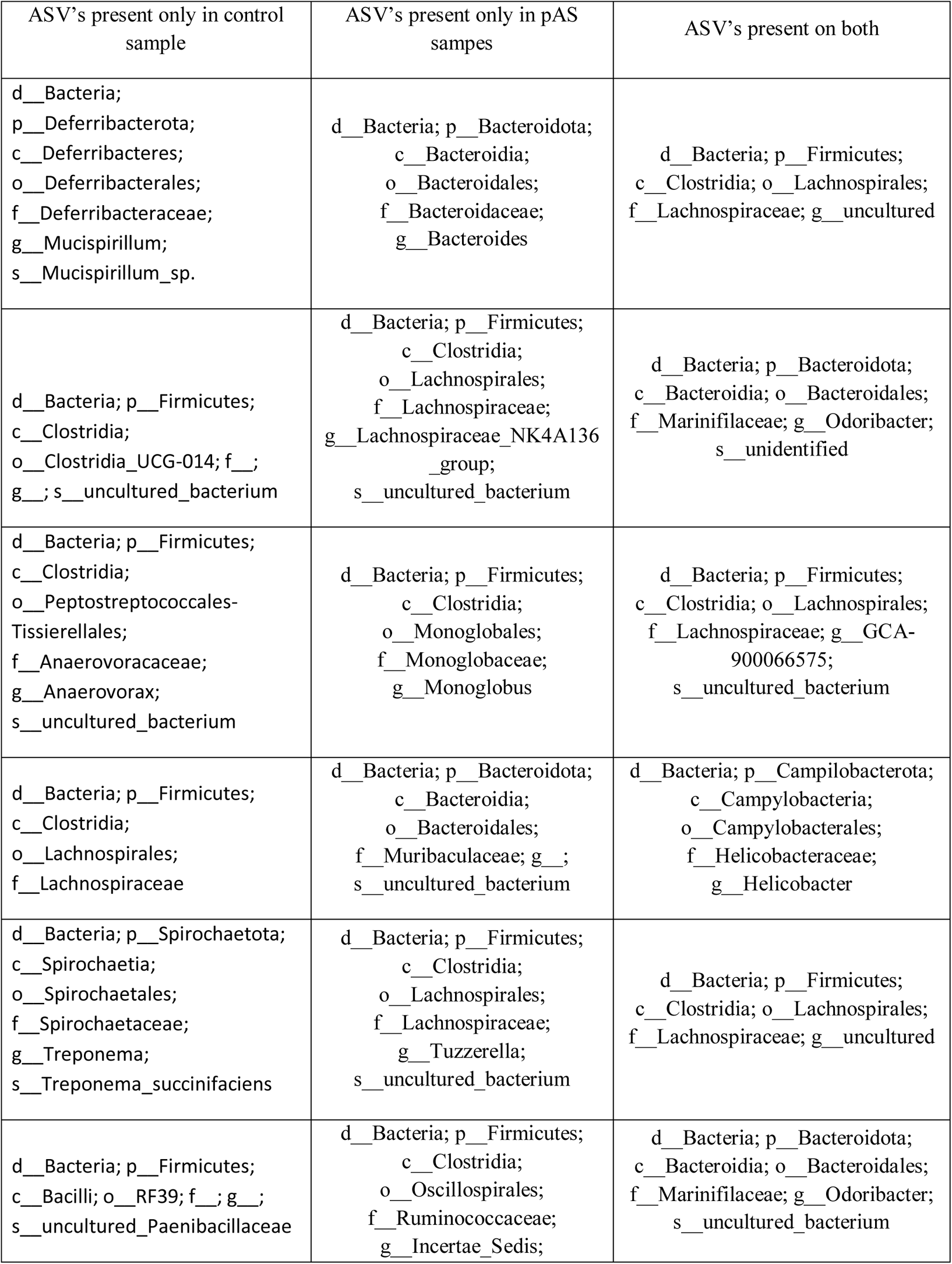
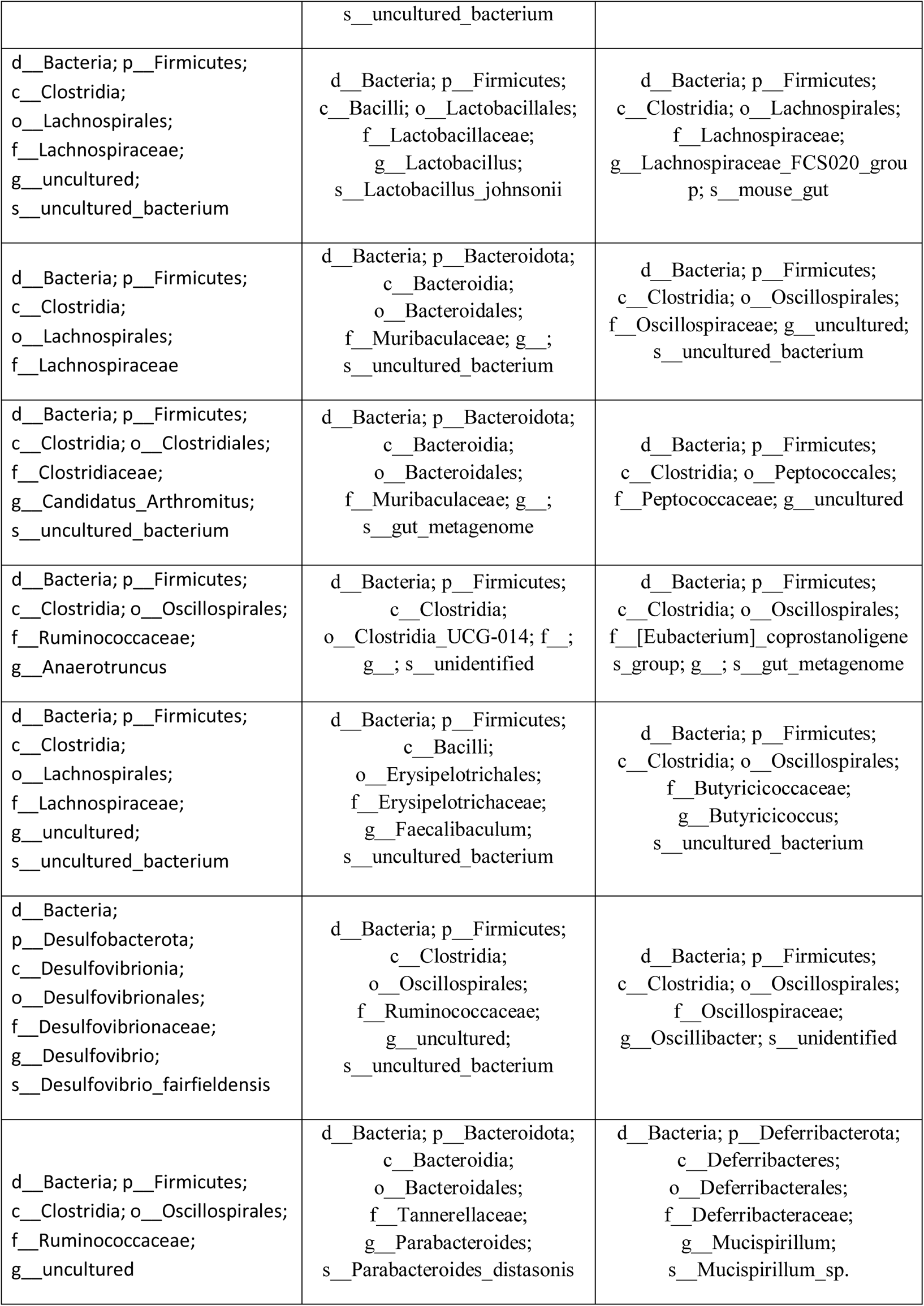
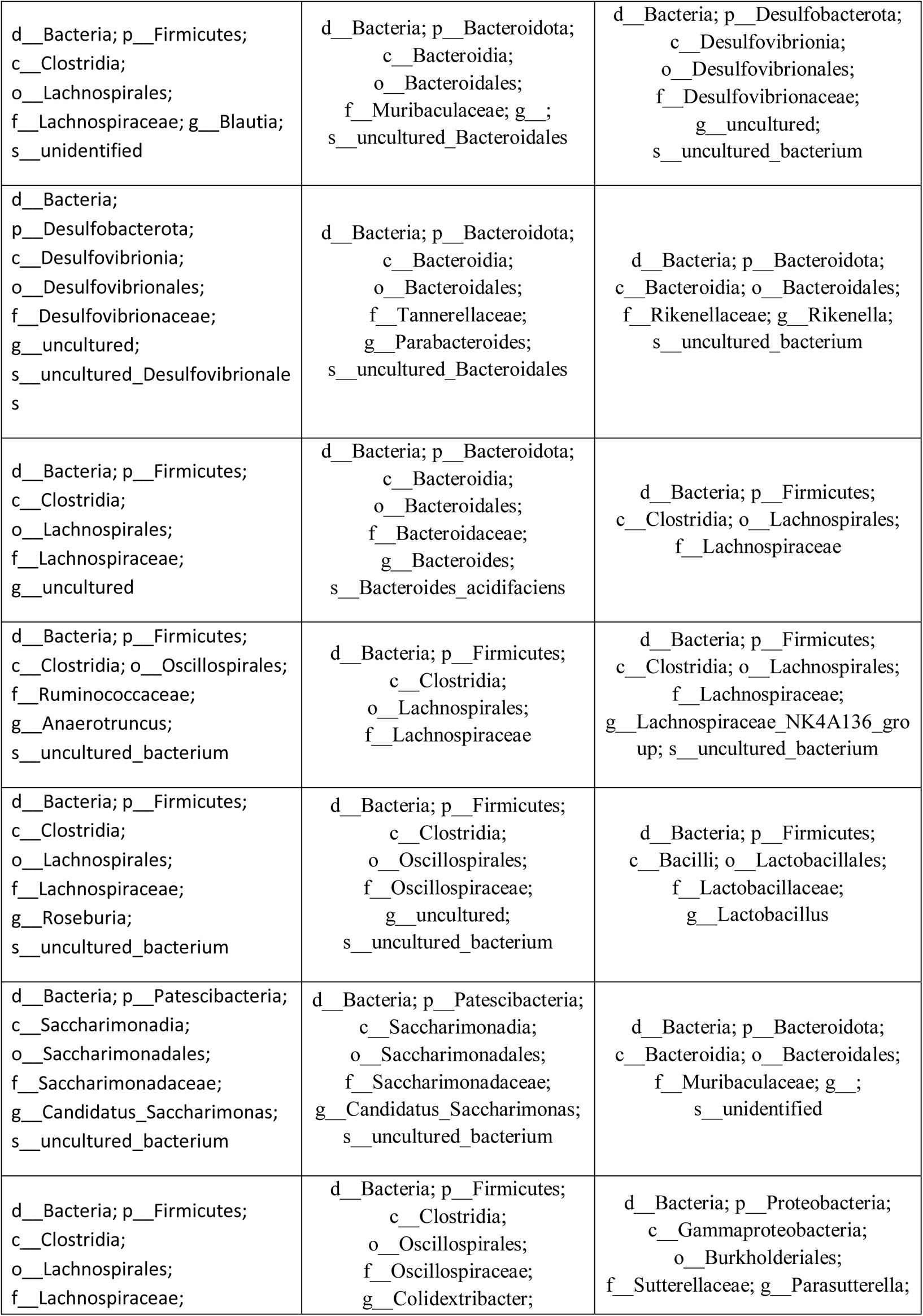
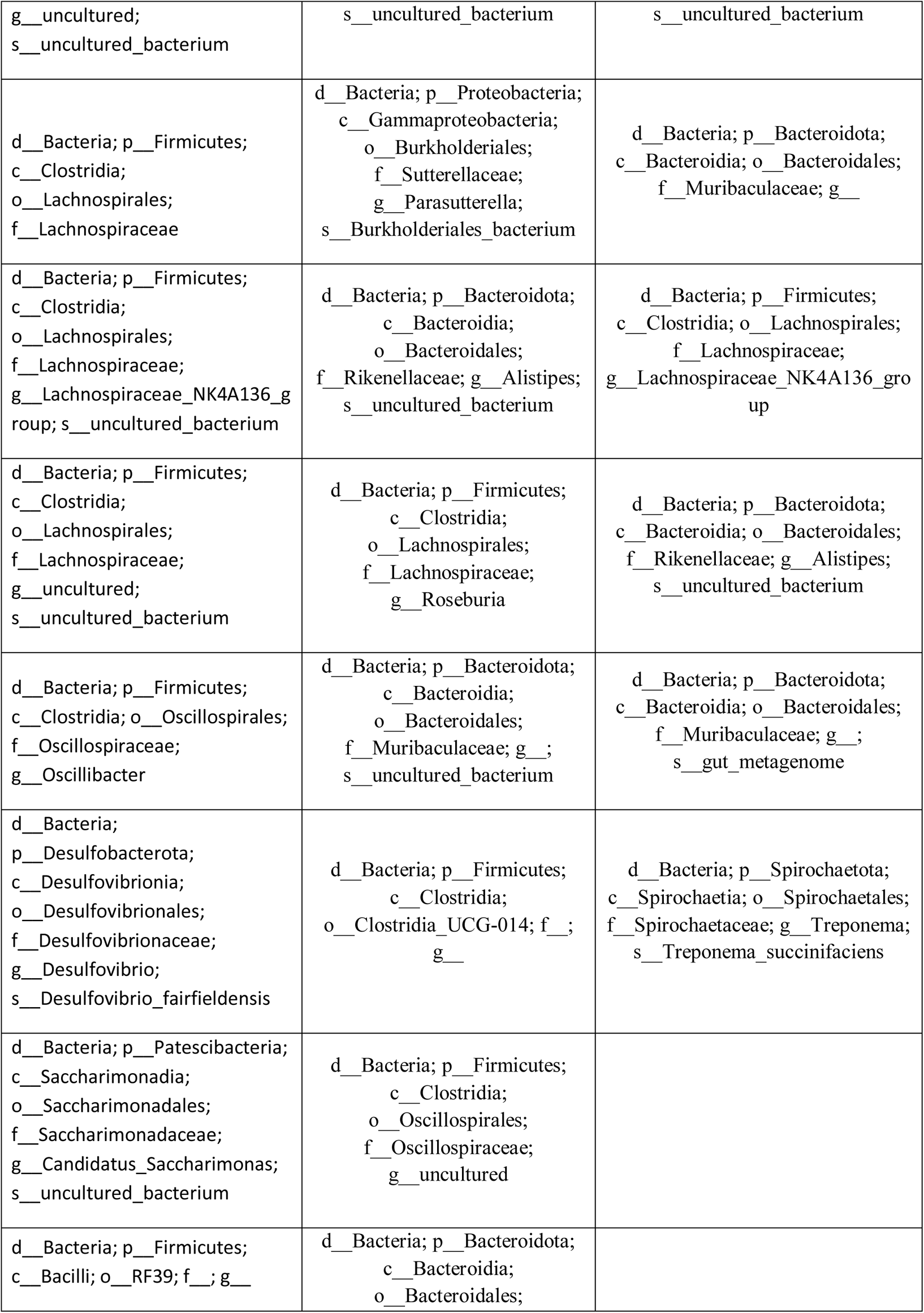
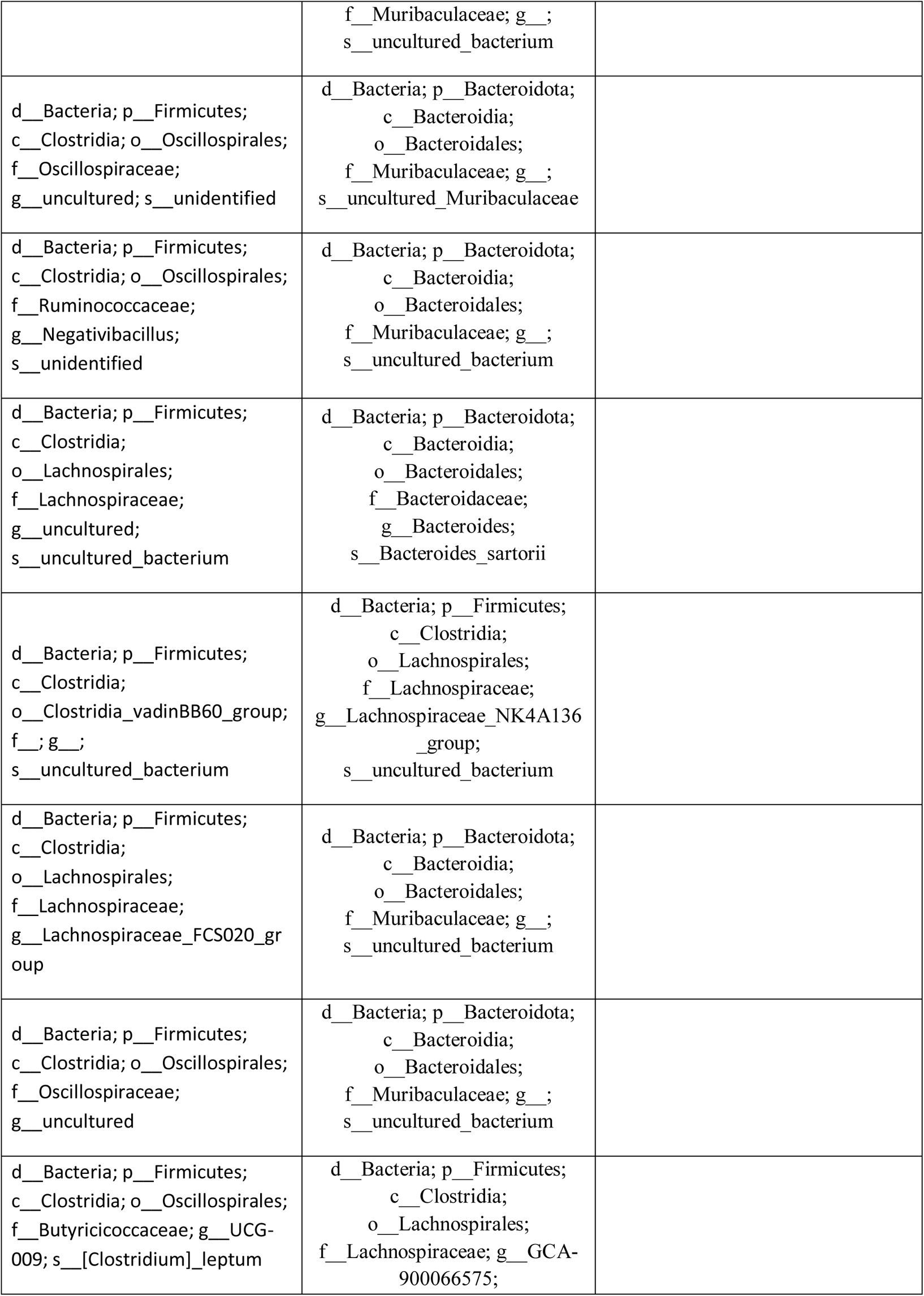
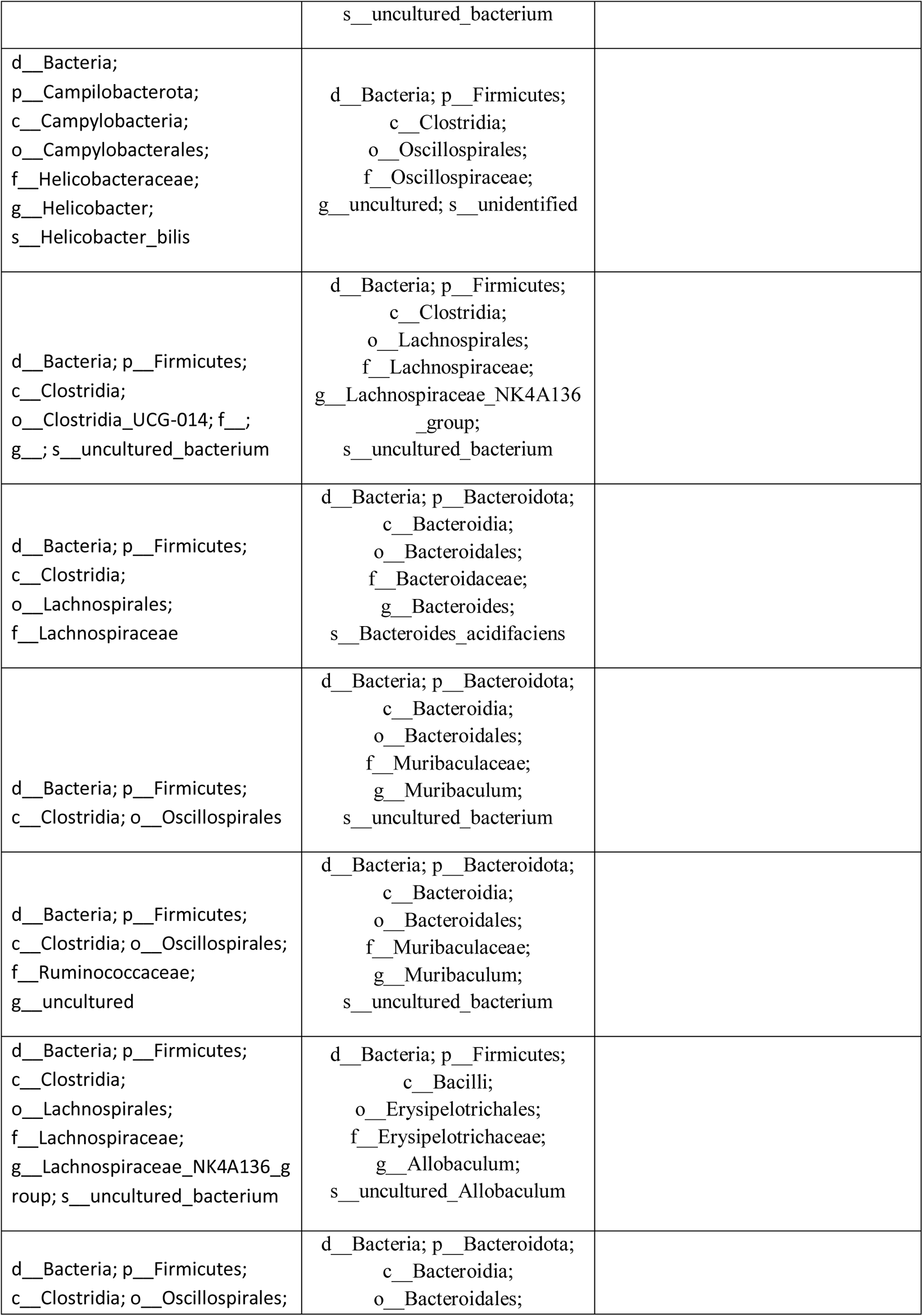
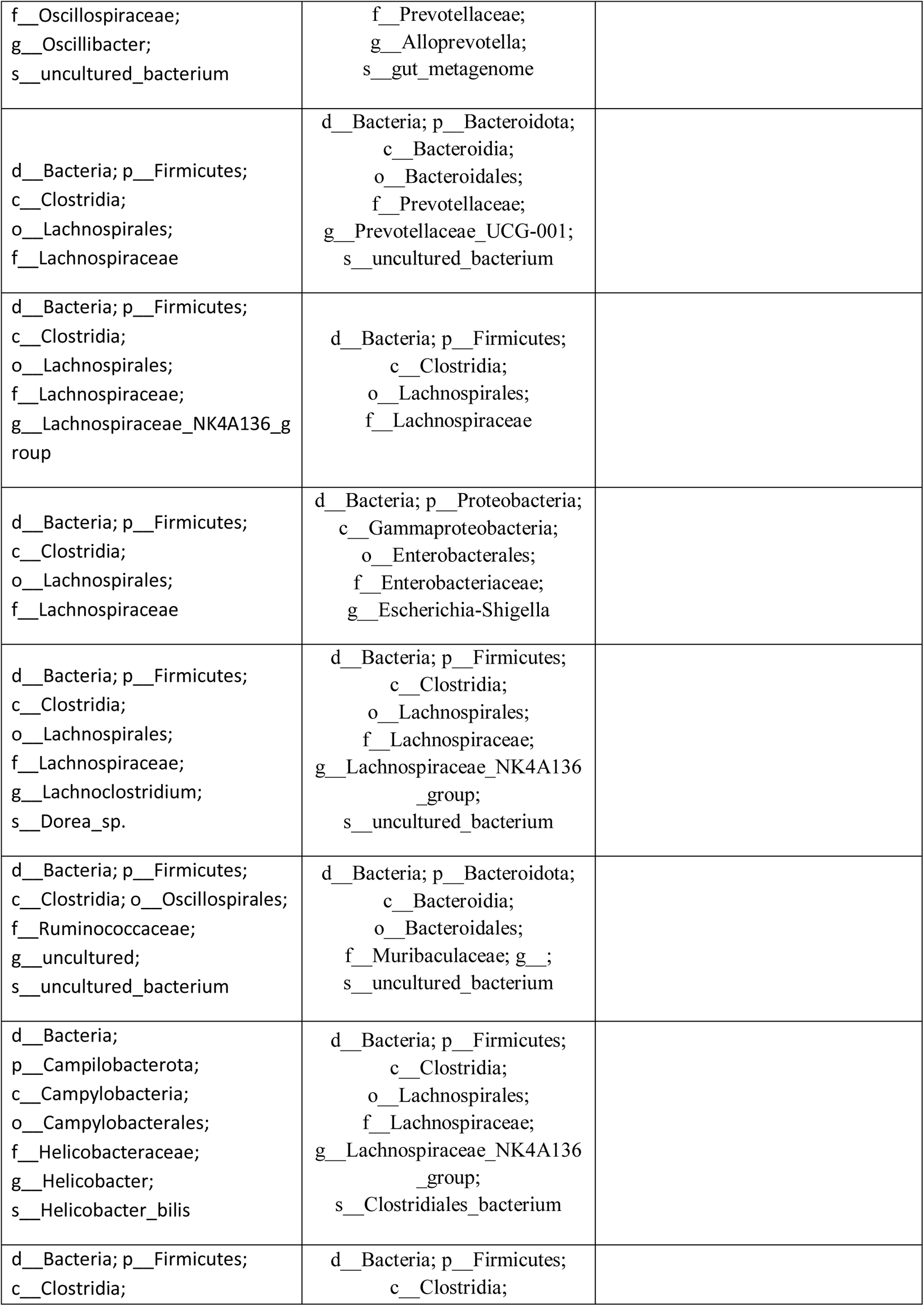
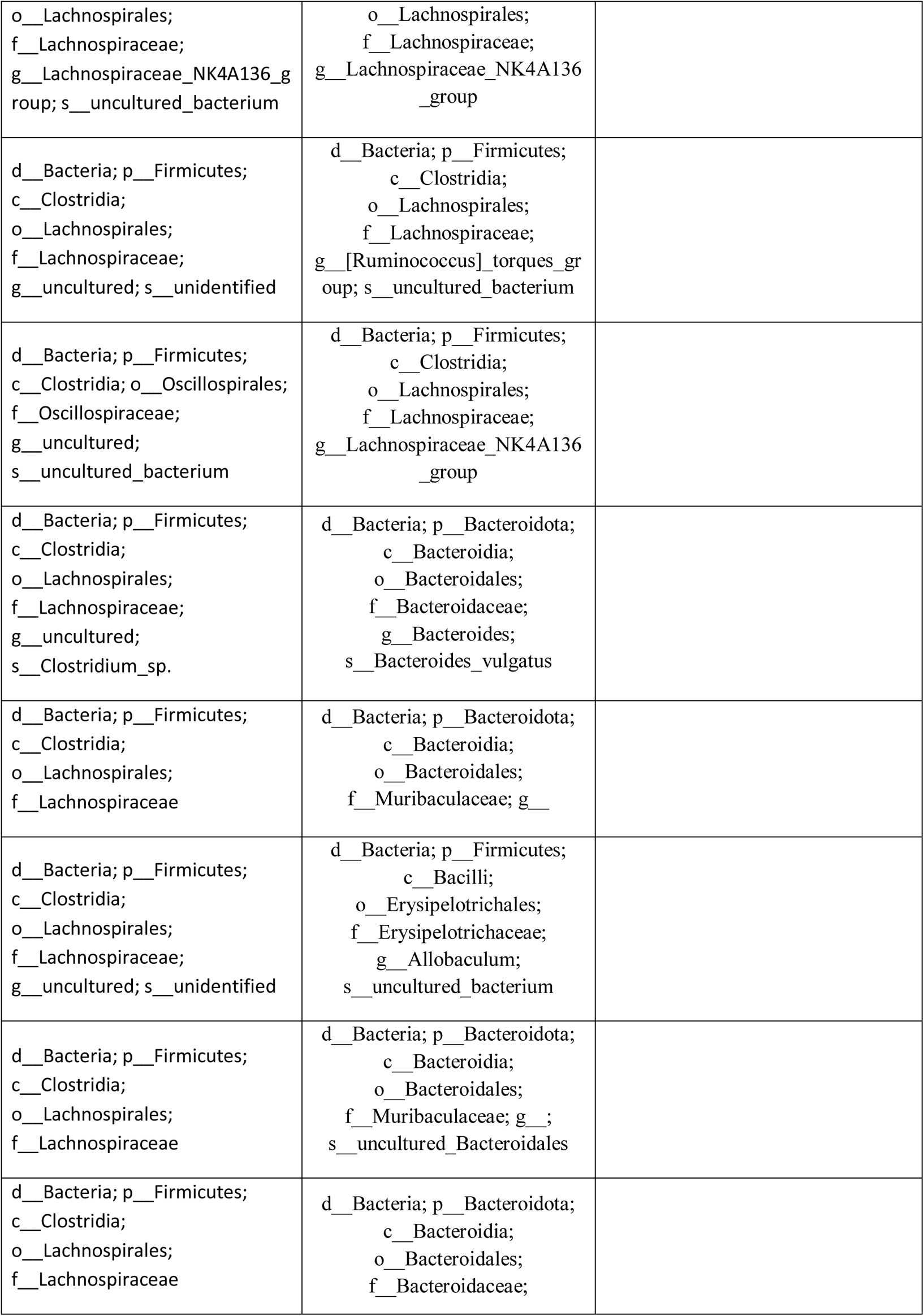
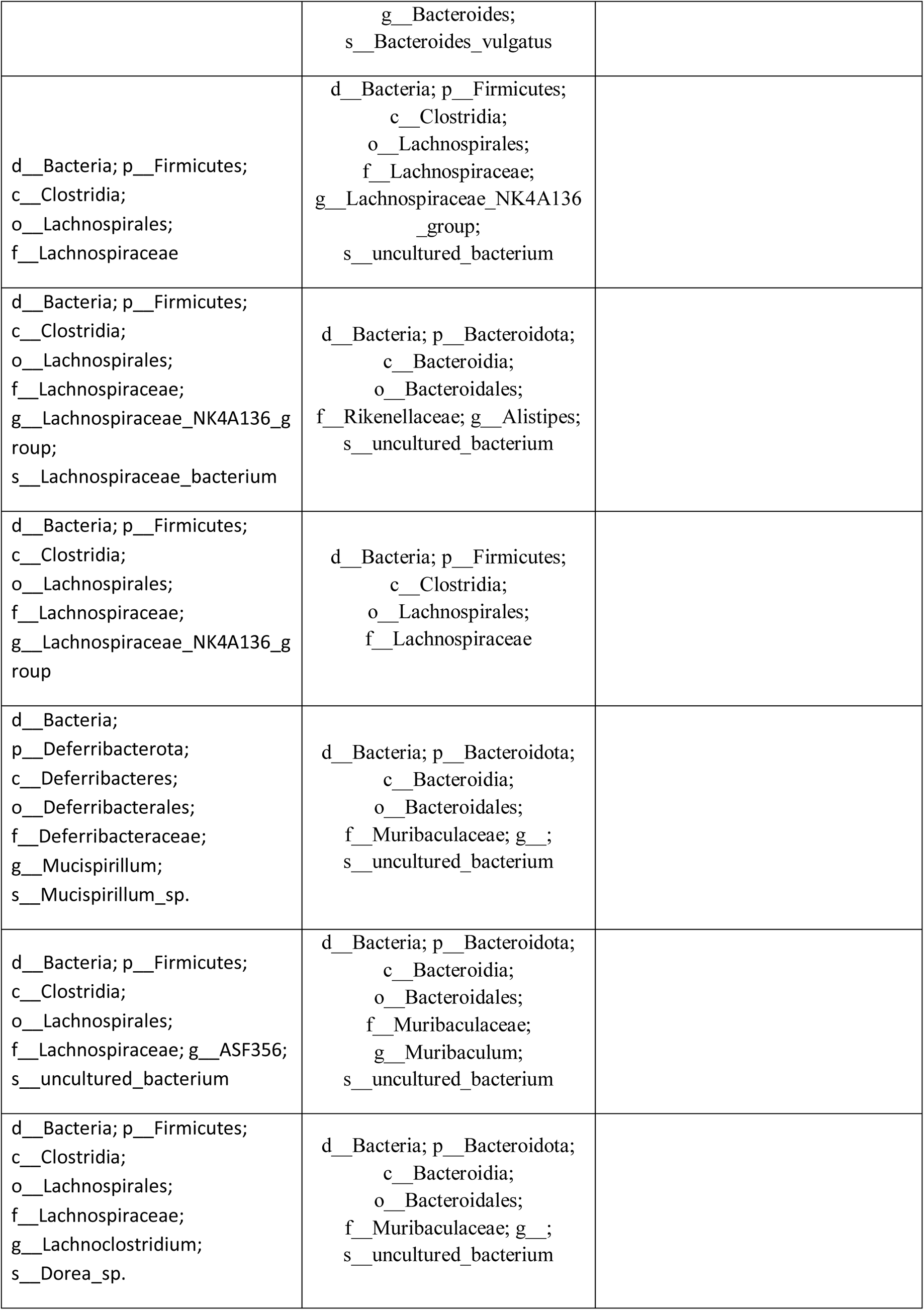
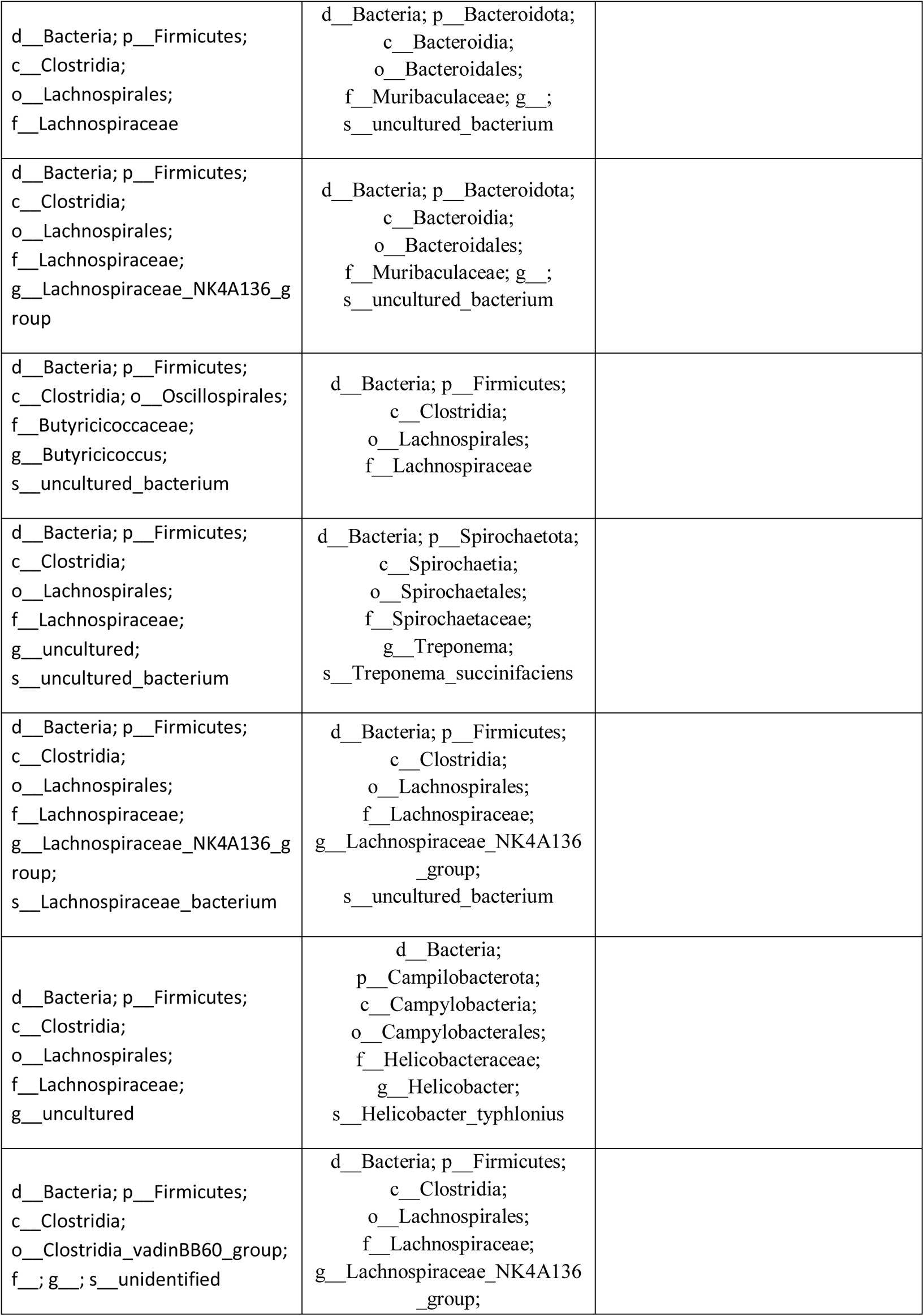
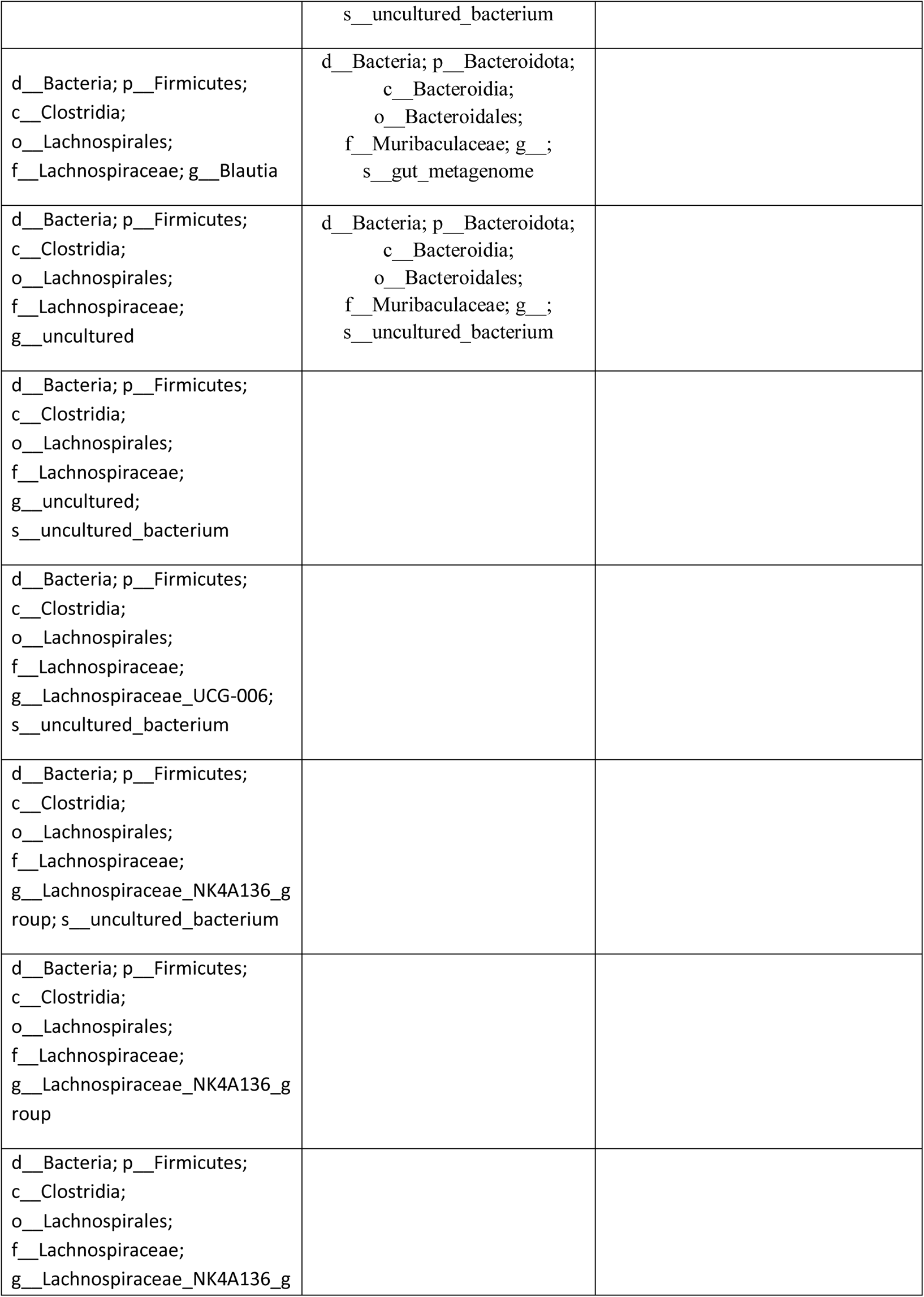
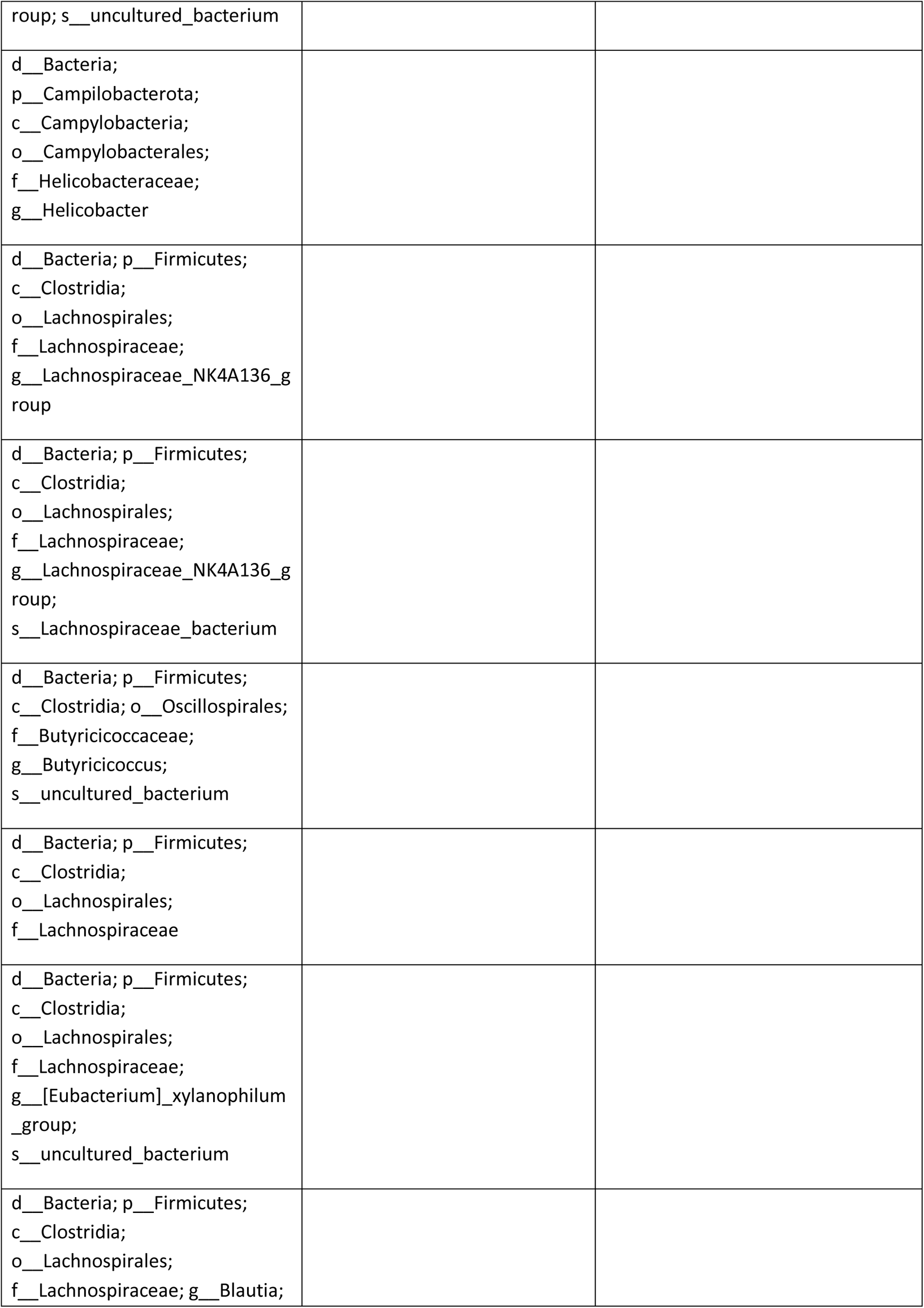
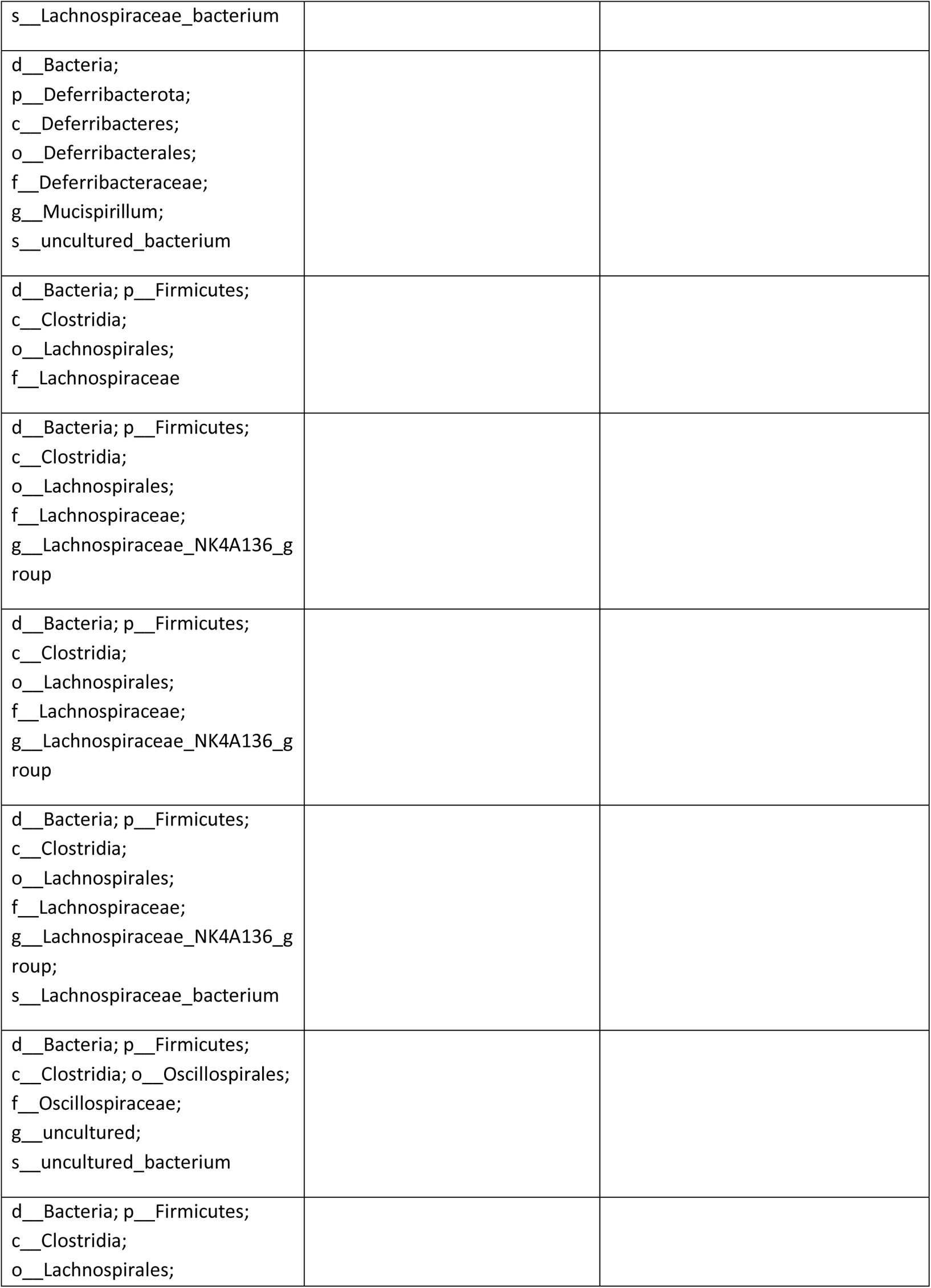
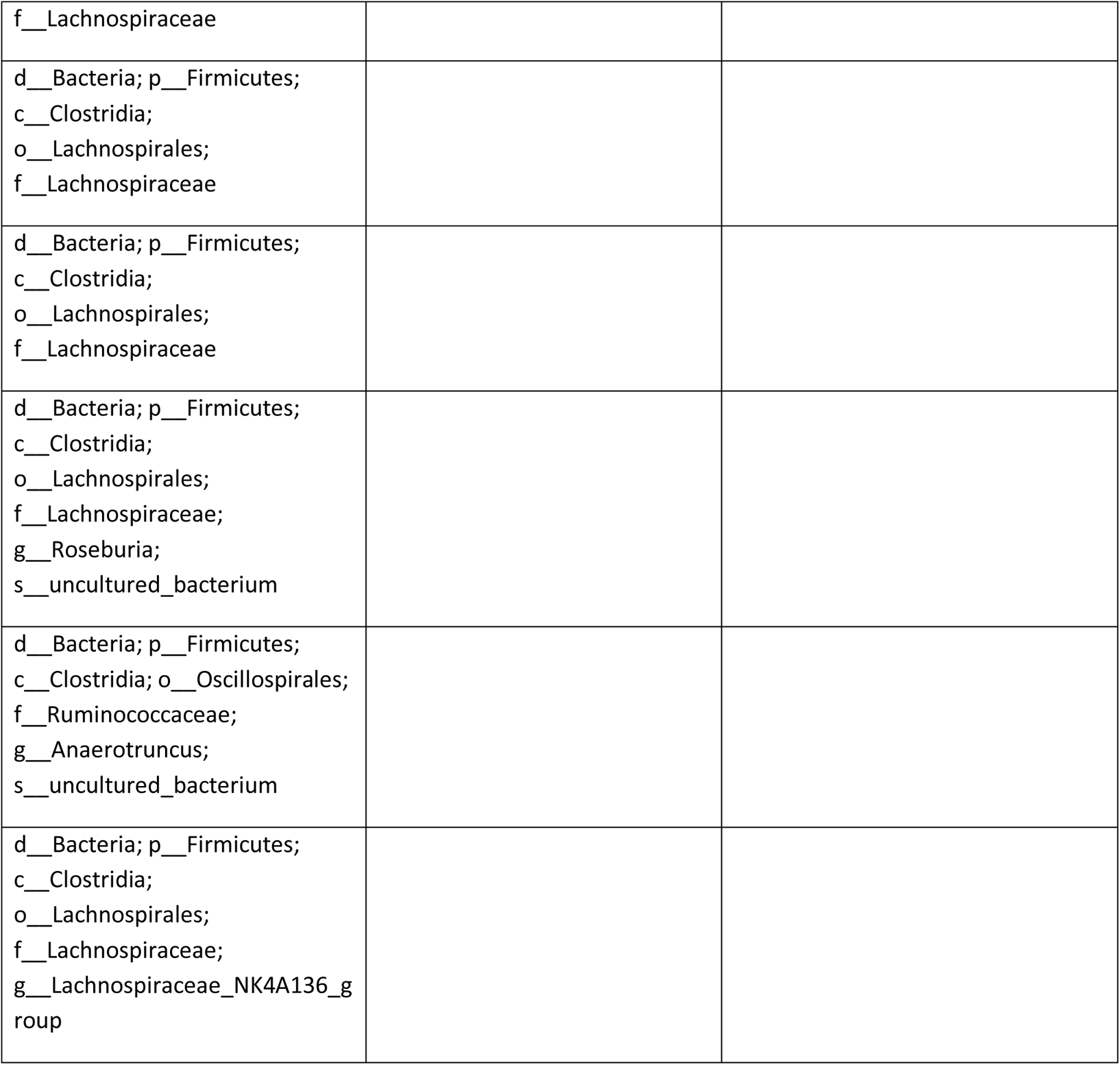
List of Genera identified in Control-mice and pAs-mice

